# Site-specific covalent labeling of large RNAs with nanoparticles empowered by expanded genetic alphabet transcription

**DOI:** 10.1101/2020.04.01.019976

**Authors:** Yan Wang, Yaoyi Chen, Yanping Hu, Xianyang Fang

## Abstract

Conjugation of RNAs with nanoparticles is of significant importance for its numerous applications in biology and medicine, which however remains challenging, especially for large ones. So far, the majority of RNA labeling rely on solid-phase chemical synthesis, which is generally limited to RNAs smaller than 100 nts. We here present an efficient and generally applicable labeling strategy for site-specific covalent conjugation of large RNAs with gold nanoparticle (AuNP) empowered by expanded genetic alphabet transcription. We synthesize an amine-derivatized TPT3 (TPT3^A^), which are site-specifically incorporated into a 97-nt 3’SL RNA and a 719-nt mini genomic RNA (DENV-mini) from Dengue virus serotype 2 (DENV2) by standard *in vitro* transcription with expanded genetic alphabet containing the A-T, G-C natural base pairs and the TPT3-NaM unnatural base pair. TPT3 modification cause minimal structural perturbations to the RNAs by small angle X-ray scattering. The purified TPT3^A^-modified RNAs are covalently conjugated with mono-*Sulfo*-NHS-Nanogold nanoparticles *via* the highly selective amine-NHS ester reaction and purified under non-denaturing conditions. We demonstrate the application of the AuNP-RNA conjugates in large RNA structural biology by an emerging molecular ruler, X-ray scattering interferometry (XSI). The inter-nanoparticle distance distributions in the 3’SL and DENV-mini RNAs derived from XSI measurements support the hypothetical model of flavivirus genome circularization, thus validate the applicability of this novel labeling strategy. The presented strategy overcomes the size constraints in conventional RNA labeling strategies, and is expected to have wide applications in large RNA structural biology and RNA nanotechnology.

**Significance Statement:** We present a site-specific labeling strategy for large RNAs by T7 transcription with expanded genetic alphabet containing TPT3-NaM unnatural base pair. The applicability of this labeling strategy is validated by X-ray scattering interferometry measurements on a 97-nt and a 719-nt RNAs. This strategy can be applicable to natural RNAs or artificial RNA nanostructures with sizes from tens up to thousands of nucleotides, or covalent conjugation of RNAs with other metal nanoparticles. The usage of a far upstream forward primer during PCR enables easy purification of RNA from DNA templates, the non-denaturing conditions for conjugation reactions and purification avoids potential large RNA misfolding. This labeling strategy expands our capability to site-specifically conjugate RNA with nanoparticles for many applications.

## Introduction

Many RNAs, including the long non-coding RNAs that are arbitrarily defined as transcripts longer than 200 nucleotides (nts) but with no or little protein coding potentials, have been found to play widespread and crucial roles in a variety of biological processes and emerge as key players in the etiology of several disease states (1). The various functions of RNAs are dictated by their propensities to form stable and complex structures (2). In light of the structure and function properties of natural RNA, a large variety of RNA nanostructures have been designed and constructed for diverse biotechnology applications such as biosensing and therapeutics over the past decade (3, 4). To elucidate and manipulate the natural and artificial RNA structures, the ability to precisely incorporate either useful labels or reactive groups to specific regions within RNA molecules is required (4, 5). These labels, conjugations are essential for further investigation and applications of RNAs as they enable the structural elucidation, visualization, localization, and biodistribution, etc (6).

Conjugation of nanoparticles (NP), particularly metal nanoparticles to biomolecules (protein, DNA, RNA) is of significant importance because of numerous applications in medicine and biology, such as in sensing, imaging, diagnosis, targeted delivery, therapeutics and structural biology (7). Due to their amenability of synthesis, unique surface, high electron density and strong optical absorption, gold nanoparticles (AuNPs) are an obvious choice in many biomedical research (8). For example, AuNPs have been used widely for labeling proteins for molecular localization in electron microscopy (9), or as markers to directly visualize the structure and conformational changes of biomolecules by atomic force microscopy (AFM) imaging (10).

Small angle X-ray scattering (SAXS) is an evolving structural technique that provides information about the shape, structure and dynamics of biomolecules under varying solution conditions (11). It has been widely used for structural analysis of RNAs (12), DNA/RNA nanostructures (13-15), or as restraints in RNA 3D structures modeling (16). Recently, it has been found that the information content of SAXS measurements can be significantly enhanced by conjugating biomolecules or DNA nanostructures with a single or a pair of AuNPs (17-20). SAXS measurements of biomolecules labeled with a single AuNP can reliably identify the label positions in the low-resolution electron density maps reconstructed from SAXS data (19). SAXS measurements of biomolecules labeled with pairs of AuNPs emerges as a new molecular ruler termed X-ray scattering interferometry (XSI) (20). XSI can provide high-resolution label-label distance distributions ranging from 50 up to 400 Å in a model-independent fashion (17, 18), thus can help to quantify ensembles of macromolecule structures and be directly related to three-dimensional structures (20). Over the last few years, XSI has been fruitfully used in nucleic acids and nucleic acid/protein complexes research (21), such as probing the structure and conformational changes of DNA origami (13), DNA or DNA-protein complexes (17, 18, 22), the conformational landscape of a complex RNA motif in response to changes in solution condition and protein binding, etc (23, 24).

The prerequisite for successful applications of XSI is to prepare biomolecules conjugated with a single and/or a pair of AuNPs at specific sites (20, 21). While labeling techniques have been extensively developed for proteins and DNAs (9, 17, 18, 22, 25, 26), nanoparticle conjugation to RNAs is more challenging, especially for large RNAs. So far, such RNAs labeling are limited to end labeling or internal labeling through solid-phase chemical synthesis (19, 21, 23, 24). The chemical synthesis method has great capability and flexibility in generating chemical diverse RNAs, but is generally limited to RNAs smaller than 100 nts. The size limits might be mitigated by combining chemical synthesis with splint-assisted enzymatic ligation, where larger RNAs are assembled from smaller pieces (5, 27), or by the complementary-addressed approaches (28, 29) where custom-designed DNA probes were utilized to direct the incorporation of reactive groups into RNAs for subsequent functional coupling. All these approaches are laborious, resulting in low yield of the final products. Also, the denaturing steps to remove the DNA splints or probes during purification are likely to prompt misfolding of RNAs (27-29), especially for large ones. Alternative approaches are therefore highly desirable for site-specific conjugation of large RNAs with nanoparticle under non-denaturing conditions.

In this work, we demonstrate an easy, efficient and generally applicable strategy for site-specific nanoparticle labeling of large RNAs empowered by expanded genetic alphabet transcription. In the past decades, three groups of Benner, Hirao and Romesberg have developed different types of unnatural base pair (UBP) that can function as a third base pair in replication, transcription, and/or translation, thus expanding the genetic alphabet (30). The expanded genetic alphabet can be used to direct the site-specific incorporation of a functionalized unnatural nucleotides into DNAs via PCR or RNAs via *in vitro* transcription catalyzed by RNA polymerases (RNAP) like T7 (31, 32). Recently, the TPT3-NaM unnatural base pair originally developed in Romesberg’s group (**Fig. 1A**) is reported to exhibit natural-like efficiency and fidelity in *in vitro* replication and transcription (33). We here synthesize an amine-derivatized TPT3 (TPT3^A^) (**Fig. 1B**), which are site-specifically incorporated into a 97-nt 3’SL RNA and a 719-nt mini genomic RNA (DENV-mini) from Dengue virus serotype 2 (DENV2) (**Fig. 2**) by standard *in vitro* transcription. The purified unnatural RNAs are covalently conjugated with mono-*Sulfo*-NHS-Nanogold nanoparticles *via* the highly selective amine-NHS ester reaction and purified under non-denaturing conditions (**Fig. 1C**). We demonstrate the application of the AuNP-RNA conjugates in large RNA structural biology by XSI measurements. The inter-nanoparticle distance distributions in the 3’SL and DENV-mini RNAs measured by XSI support the hypothetical model of flavivirus genome circularization, thus validate the applicability of this novel labeling strategy. The presented strategy overcomes the size constraints in conventional RNA labeling strategies, and is expected to have wide applications in large RNA structural biology and RNA nanotechnology.

**Fig. 1.**
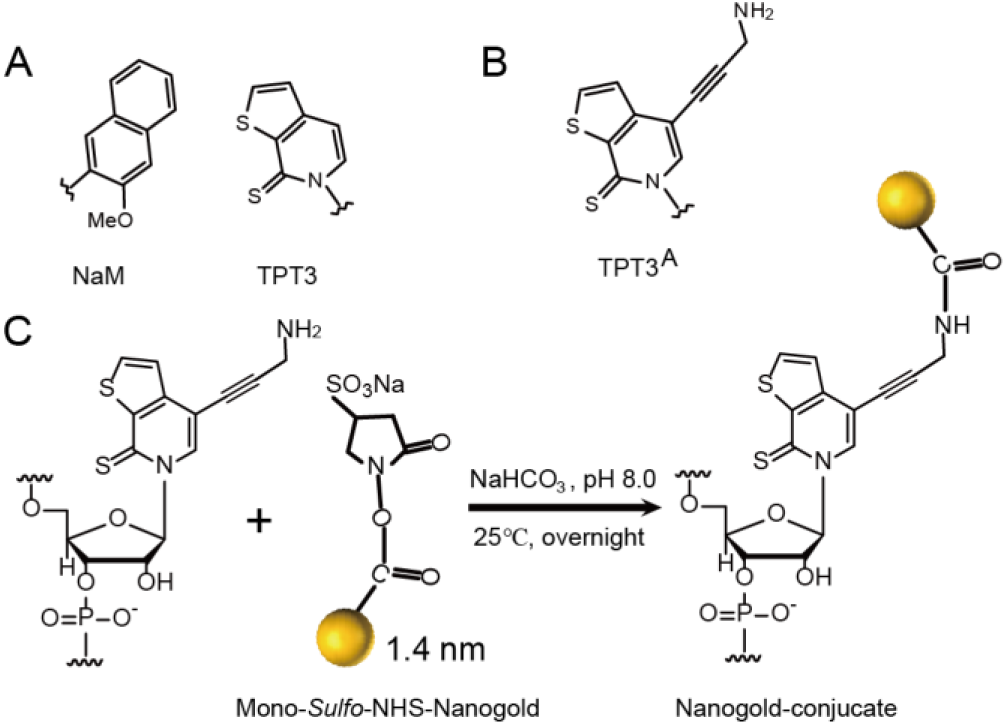
The TPT3-NaM unnatural base pair (UBP) for genetic alphabet expansion. (*A*) The parental TPT3-NaM UBP. (*B*) Chemical structure of the amine-derivatized TPT3 (TPT3^A^). (*C*) Schematic diagram for covalent conjugation of TPT3^A^-modified RNAs with mono-*sulfo*-NHS-Nanogold via amine-NHS ester reaction.

**Fig. 2.**
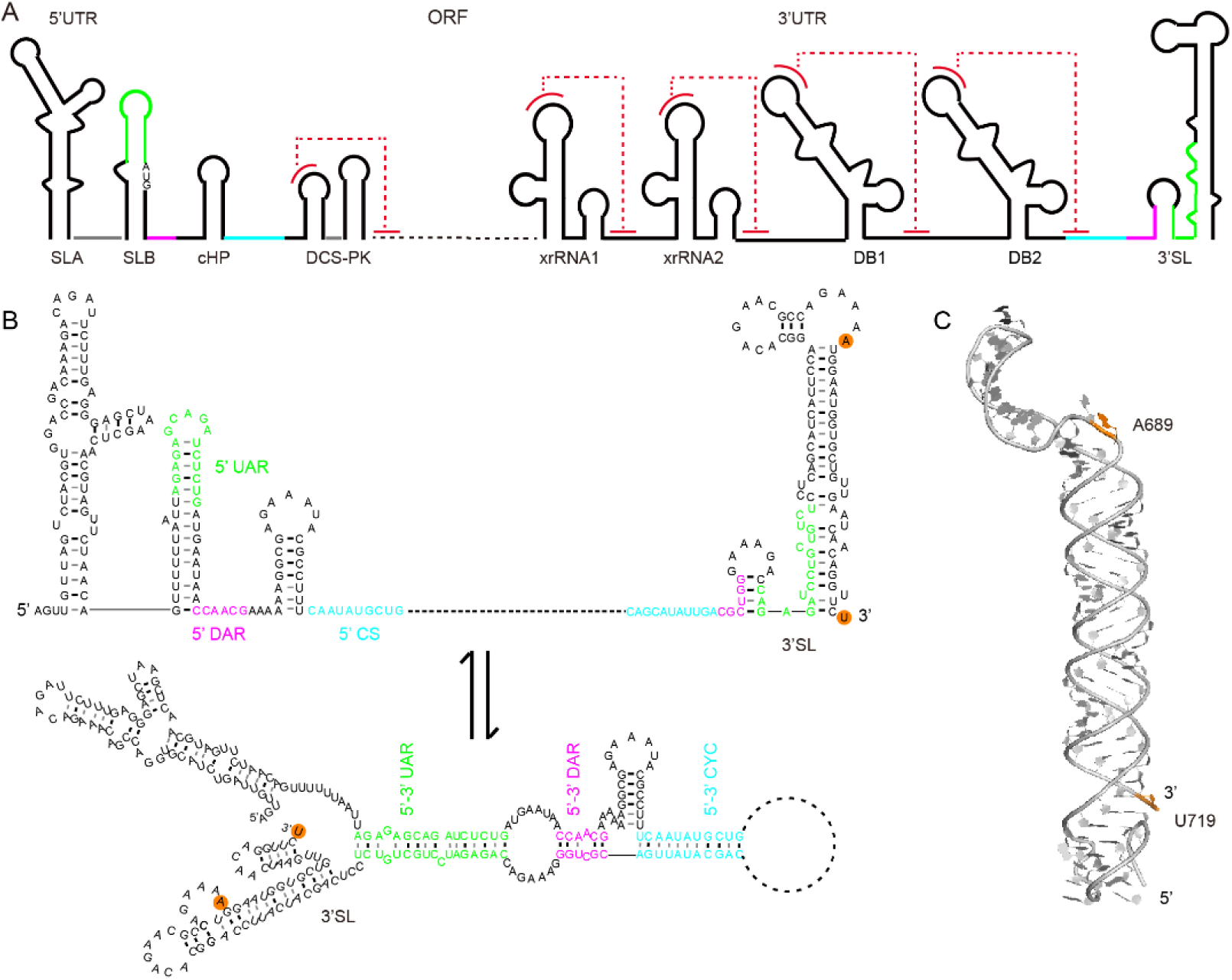
Conformational changes of the 3’SL RNA element of Dengue virus 2 (DENV2) upon genome cyclization. (*A*) The single-stranded positive sense genomic RNA of DENV2 consists of an open reading frame (ORF) flanked with many highly structured RNA elements, such as the SLA, SLB, cHP and DCS-PK elements located in the 5’-untranslated regions (UTR) and the capsid protein coding regions, the xrRNA1, xrRNA2, DB1, DB2 and 3’SL elements located within the 3’ UTR. Pseudoknots formation in these elements are colored by dashed lines in red. (*B*) Three sets of long-range RNA-RNA interactions: 5’-3’ upstream of AUG region (UAR) (green), 5’-3’ downstream of AUG region (DAR) (magenta) and 5’-3’ cyclization sequence (CS) (cyan) mediate the linear-to-circular conformational transition of the genomic RNA upon genome cyclization, resulting in significant structural changes in RNA elements, such as the SLB and 3’SL. The labeling sites corresponding to A689 and U719 in DENV-mini are indicated with orange background. (*C*) Atomic model of the 97-nt 3’SL element by SAXS and computation, which forms an extended conformation co-axially stacked by the small hairpin (sHP) and the long stem loop (SL)(39). The distance between N9 of A689 and N1 of U719 is measured as 83.3 Å in the atomic model.

## Results

### Chemical Synthesis of Unnatural Nucleotides

To implement *in vitro* replication and transcription by the expanded genetic alphabet containing TPT3-NaM (**Fig. 1A**), the deoxyribonucleotide phosphoramidites, triphosphorylated deoxy- and ribonucleotides of TPT3 and NaM (dTPT3-CEP, dNaM-CEP, dTPT3TP, dNaMTP, rTPT3TP, rNaMTP) were synthesized according to the literature procedures (31, 33, 34). Since extremely uniform 1.4 nm gold nanoparticle (Nanogold) with a single sulfo-N-hydroxysuccinimide ester (Mono-sulfo-NHS) that can react with primary amines is commercially available, we synthesized amine-modified triphosphorylated ribonucleotides of TPT3 (rTPT3^A^TP) (**Fig. 1B**) as described in the ***SI Appendix, S1***. rTPT3^A^TP is expected to be incorporated into RNA transcripts by expanded genetic alphabet transcription at specific sites, and the TPT3^A^-modified RNAs are expected to allow stable, covalent conjugation of the Nanogold via amine-NHS ester reaction in solution (**Fig. 1C**).

### Selection of Labeling Sites

Dengue virus is an important human pathogen featuring a single-stranded positive-sense RNA genome which contains an open reading frame (ORF) flanked by highly structured untranslated regions (UTR) (**Fig. 2A**). During infection, the DENV genome serves as mRNA for translation, template for RNA synthesis and substrate for encapsidation and is predicted to undergo conformational transition between the linear and circular forms to fulfill its diverse function (35, 36). The circular conformation is stabilized by long-range RNA-RNA interactions mediated by the inverted complementary sequences at both the 5’ and 3’ ends of the genome (**Fig. 2B**) (37), resulting in conformational changes in several conserved RNA structures. One example is the 3’SL structure, which consists of a small hairpin (sHP) followed by a large stem loop (SL) and is highly conserved among all flavivirus genomes (**Fig. 2B**) (38). Upon genome circularization, the 3’SL RNA element is expected to open the large stem and release the last nucleotide of the genome, thus adopts the circular conformation that has a large stem loop ahead of a small one (**Fig. 2B**) (35, 36), acting as template RNA to initiate replication. However, the tertiary conformational changes have not been validated before.

The 97-nt 3’SL RNA alone (***SI Appendix*, Fig. S1A**) was recently confirmed to have an extended rod-like structure in solution by SAXS (**Fig. 2C**) (39), thus representing the linear conformation of the genomic RNA. The 719-nt DENV-mini RNA, which consists of all the elements in 5’ UTR, 3’UTR and part of the capsid coding sequence of the genomic RNA, is a minimal and efficient template for translation and minus strand RNA synthesis *in vitro* (40). The secondary structure of DENV-mini (***SI Appendix*, Fig. S1B**) was recently probed by SHAPE probing, which verified the long-range RNA-RNA interactions between the 5’ and 3’ terminal regions that mediates genome circularization (41). Thus, the smaller DENV-mini RNA can resemble the circular conformation of the full length genomic RNA of DENV2, which is generally larger than 11 kb. To study the conformational changes of the 3’SL upon genome circularization by XSI, two sites in both 3’SL and DENV-mini corresponding to A689 and U719 (referred to DENV-mini) were selected to be labeled with TPT3^A^ for Nanogold conjugation. Based on the secondary structures, A689 and U719 are located at the internal loop and 3’-terminal regions, respectively, in both 3’SL and DENV-mini (***SI Appendix*, Fig. S1**), incorporation of TPT3^A^ at these sites are expected to avoid significant structural perturbations and allow for conformational changes to be monitored.

### Preparation of DNA Templates containing UBPs at Specific Sites

Previously, it was reported that dNaM in the template strand of double-stranded DNA (dsDNA) template can direct the incorporation of its partner into RNA more efficiently and selectively (42). To prepare single- and/or double-site TPT3 or TPT3^A^ labeled RNAs by expanded genetic alphabet transcription, we generate dsDNA templates containing one or two dNaMs in the template strand at specific sites by PCR reactions. Two plasmids coding for 3’SL and DENV-mini RNAs with an upstream T7 promoter, respectively, are total gene synthesized as PCR templates (***SI Appendix*, Table S1**). To introduce the UBPs into the dsDNA templates, the UBPs are first incorporated into the single-stranded DNA primers using standard solid-phase chemical synthesis with the phosphoramidites of dNaM or dTPT3 (**Fig. 3A**). As the two labeling sites (A689 and U719) are close to the 3’-end of 3’SL, only a pair of forward and reverse primers are needed for generating the dsDNA templates in one PCR reaction for each constructs. In case of preparation of dsDNA templates with two distant labeling sites, two pairs of primers can be synthesized and an overlap extension PCR protocol can be employed (43, 44). A common forward primer pMVF along with three reverse primers containing one or two dNaMs at sites corresponding to A689 and/or U719 were initially synthesized (***SI Appendix*, Table S2**). The forward primer pMVF was designed to target a common sequence 390 base-pair (bp) upstream of the T7 promoter in the plasmids (**Fig. 3A**), such a design results in dsDNA templates significantly larger than the RNA transcripts, enabling subsequent efficient and easy RNA purification (see below). Directed by the respective pair of the native forward and unnatural reverse primers, 6 letter PCR reactions were performed with a mixture of dNTPs of A, T, G, C, NaM and TPT3. The PCR amplifications produce dsDNA templates containing one or two dNaMs in the template strand at sites corresponding to A689 or/and U719 for both 3’SL and DENV-mini RNAs, with lengths of 504 and 1128 bps, respectively (***SI Appendix*, Fig. S2**). Since the TPT3-NaM UBP has a natural base pair-like efficiency and fidelity in *in vitro* replication and the dNaMs are firstly introduced into the reverse primers, no mutagenesis in the template strand of dsDNA templates is expected during PCR amplification. The PCR products were directly used for subsequent transcription without further purification.

**Fig. 3.**
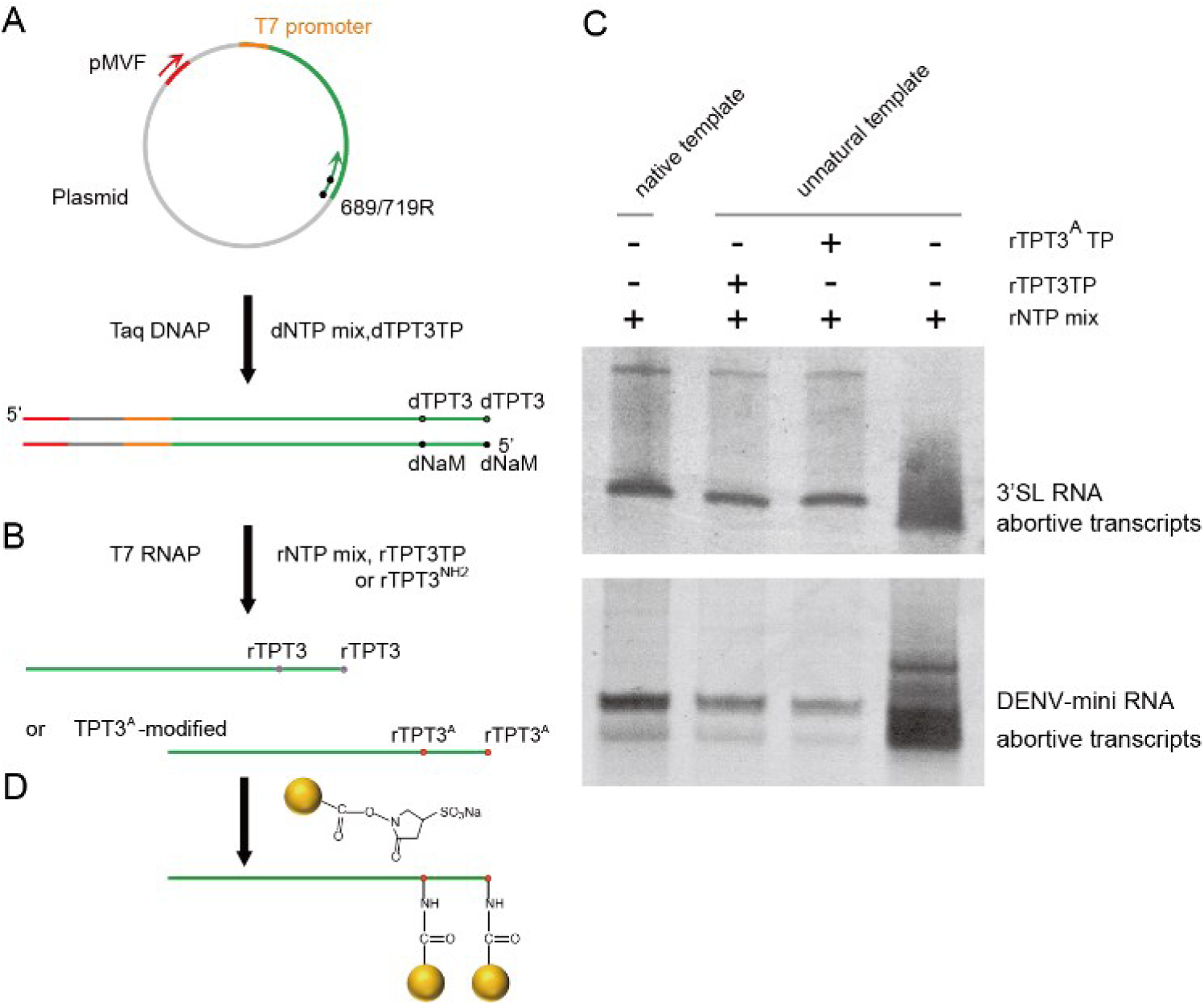
A general strategy for site-specific Nanogold labeling of large RNAs using an expanded genetic alphabet containing TPT3-NaM. (*A*) The UBPs were first incorporated into the reverse ssDNA primers targeting the plasmids by solid-phase chemical synthesis with the phosphoramidites of dNaM or dTPT3. Directed by an upstream forward primer and the respective reverse primers, dsDNA templates containing UBPs at specific sites were amplified by PCR. (*B*) rTPT3 or rTPT3^A^ can be incorporated into RNAs by *in vitro* transcription with dsDNA templates containing UBPs (dNaM in the template strand) using the six-letter expanded genetic alphabet. (*C*) 10 % (for 3’SL) or 6 % (for DENV-mini) native PAGE analysis of the *in vitro* transcripts from native DNA templates or unnatural DNA templates (A689TPT3). The transcription of native DNA template (*Left*) using rNTP mix resulted in one single band for 3’SL (top) and DENV-mini (down). The transcription of unnatural DNA template (*Right*) in the absence of unnatural nucleotides resulted in smeared bands. The transcription of unnatural DNA template (*Middle*) in the presence of rTPT3 or rTPT3^A^ resulted in dense bands. (*D*) Mono-*Sulfo*-NHS-Nanogold labels are coupled to purified TPT3^A^-modified RNAs *via* amine-NHS ester reaction.

### Site-specific Incorporation of rTPT3 or rTPT3^A^ into RNAs

Although the *in vivo* replication and transcription of TPT3-NaM UBP have been validated recently in an *Escherichia coli* and an eukaryotic yeast semi-synthetic organisms by Romesberg group (45, 46), the ribonucleotides of TPT3 and TPT3^A^ have not been tested in *in vitro* transcription with T7 RNA polymerase. The dsDNA templates containing dNaMs in the template strand for 3’SL and DENV-mini are used to explore site-specific incorporation of UBP into RNAs.

We first characterized the ability of T7 RNA polymerase to incorporate rTPT3 and rTPT3^A^ into the 97-nts 3’SL RNA (**Fig. 3B**). The transcription of a DNA template containing one dNaM at site corresponding to A689 of 3’ SL RNA was first examined with the natural ribotriphosphates and either rTPT3 or rTPT3^A^. Under the conditions employed (***SI Appendix*, Table S4**), virtually no clear full-length product but smeared bands were observed in the absence of unnatural triphosphate, this shows that rTPT3 or rTPT3^A^ is essential for correct transcription of DNA templates containing dNaM. On the contrary, addition of either rTPT3 or rTPT3^A^ resulted in efficient production of the full-length transcription product (**Fig. 3C**). These data suggest that enzymatic incorporation of both rTPT3 and rTPT3^A^ into RNA is feasible via *in vitro* T7 transcription. We further confirmed the selective incorporation of rTPT3 or rTPT3^A^ into the 719-nt DENV-mini RNA using its dsDNA template containing dNaM at site corresponding to A689 (**Fig. 3C**).

For XSI data acquisition, usually a sample quartet including the unmodified, the two orthogonal one site- and the one double-modified constructs and the free Nanogold sample are prepared (20, 21). We then prepared the A_689_TPT3/A_719_TPT3- (for the unmodified), A_689_TPT3^A^- (for one single-site Nanogold labeling), A_719_TPT3^A^- (for another single-site Nanogold labeling), A_689_TPT3^A^/A_719_TPT3^A^- (for double-site Nanogold labeling) modified RNAs of 3’SL and DENV-mini, respectively in the presence of natural ribotriphosphates (A, U, G, C) mix and TPT3 or TPT3^A^. As the dsDNA templates are much larger than the RNA transcripts, each of the RNA transcripts is well resolved from the dsDNA template in the native PAGE gel, digestion of dsDNA templates is not needed prior to native purification. The transcription supernatants are applied to the Superdex 75 (for 3’SL) or 200 (for DENV-mini) gel filtration columns, the RNAs are directly purified from the dsDNA templates, the abortive transcripts and the excess rNTPs by Size-exclusion chromatography (SEC) (***SI Appendix*, Fig. S3A**).

### Effects of TPT3 and TPT3^A^ Modification on RNA Structures

Previously, it was reported that gold nanocrystal labels do not substantially perturb RNA structures (24). We therefore investigate the influence of the incorporation of TPT3 or TPT3^A^ on RNA structures prior to Nanogold labeling by SAXS. The scattering profiles, with scattering intensity I(*q*) plotted against momentum transfer *q*, along with PDDFs (pair distance distribution function) transformed from the scattering profiles for all the UBP-modified RNAs and the natural RNAs of 3’SL and DENV-mini are shown in **Fig. 4**. The Guinier regions of all the scattering profiles are linear (**Figs. 4A and 4B**), indicating that all the RNA samples are monodispersed and homogeneous in solution. Comparing the scattering profiles and PDDFs of the TPT3- or TPT3^A^-modified RNAs with the natural RNAs of the 3’SL and DENV-mini, respectively, we concluded that TPT3 or TPT3^A^ modification has little effects on the overall structures of the respective RNAs. The overall structural parameters, including the radius of gyration *R*_g_ calculated from the slops of Guinier fitting, *R*_g_ and the maximum dimension *D*_max_ from PDDF functions, as well as molecular weights derived from volume-of-correlation (*V*_c_) (47), are summarized in ***SI Appendix*, Table S5**, among which the molecular weights calculated from SAXS data are consistent with those predicted from sequences, indicating that all the RNA constructs are monomeric in solution. The slight variations of the *R*_g_ and *D*_max_ upon UBP modification further proved the absence of significant structural changes.

**Fig. 4.**
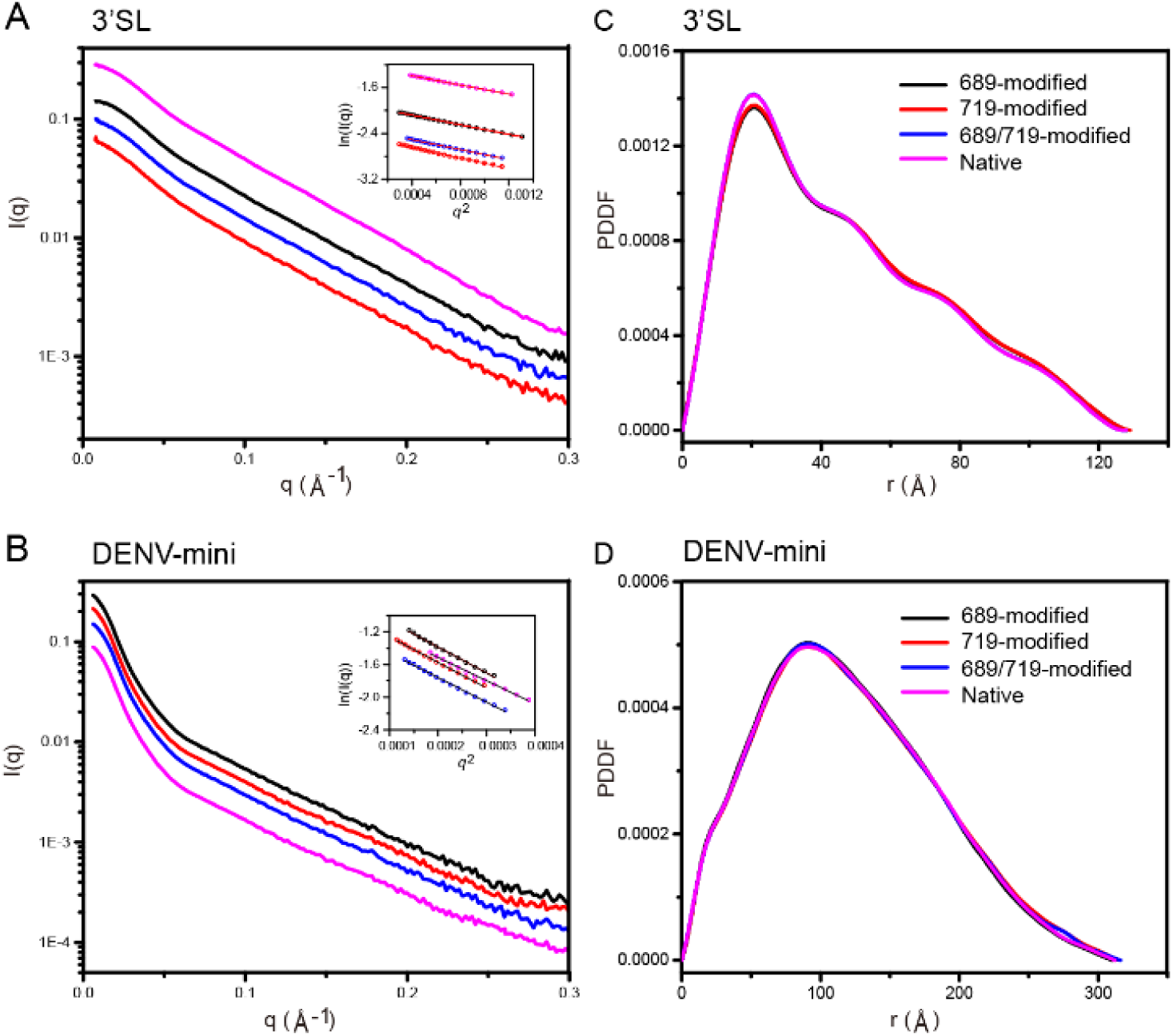
Effects of TPT3-incorporation on the structures of 3’SL and DENV-mini RNAs by SAXS. (*A***-** *B*) Experimental scattering profiles of native (magenta), 689-modified (black), 719-modified (red) and 689/719 double-modified (blue) RNAs for 3’SL (**A**) and DENV-mini (**B**). The insets in A and B are the respective linear fittings of the Guinier plotting. (*C***-***D*) Overlay of the respective PDDFs of native (magenta), 689-modified (black), 719-modified (red) and 689/719 double-modified (blue) RNAs for 3’SL (**C**) and DENV-mini (**D**).

### Site-specific Nanogold labeling of RNAs

Purified unnatural 3’SL and DENV-mini RNAs containing one TPT3^A^ at sites of 689 or 719, or two TPT3^A^ at both sites of 689 and 719 were subjected to covalent conjugation with a two-to three-fold excess of the 1.4 nm Mono-*Sulfo*-NHS-Nanogold from Nanoprobe (http://www.nanoprobes.com/), resulting in stable RNA conjugates with one site- or double-labeled Nanogold (**Fig. 3D**). Reaction of primary amine with NHS esters at physiological pH in aqueous environment is fast and highly selective, and cross-linking by this reaction is one of the most commonly used methods in RNA labeling (5). Freshly prepared Mono-*Sulfo*-NHS-Nanogold was immediately reacted with TPT3^A^- modified RNA samples in order to avoid hydrolysis of the NHS ester. The pH was maintained between 7 and 9 in 0.1 M NaHCO_3_ since the hydrolysis of NHS ester is pH dependent. The RNA conjugates were further purified from the excess Nanogold by SEC. As shown in **Fig. 5A**, the Nanogold-conjugated DENV-mini RNA can be directly purified from the excess free Nanogold with the Superdex200 gel filtration column, which shows good separation between peaks in the elution profile. However, the elution peak of Nanogold-conjugated 3’SL RNA overlaps with that of free Nanogold at both Superdex 75 and 200 gel filtration columns, the 3’SL-Nanogold conjugates were therefore purified by Anion-exchange chromatography (**Fig. 5B**). The native purification protocols for Nanogold-labeled 3’SL and DENV-mini RNAs avoid the denaturing steps used in other labeling methods (27-29), which help the RNAs preserve conformations during co-transcriptional folding.

**Fig. 5.**
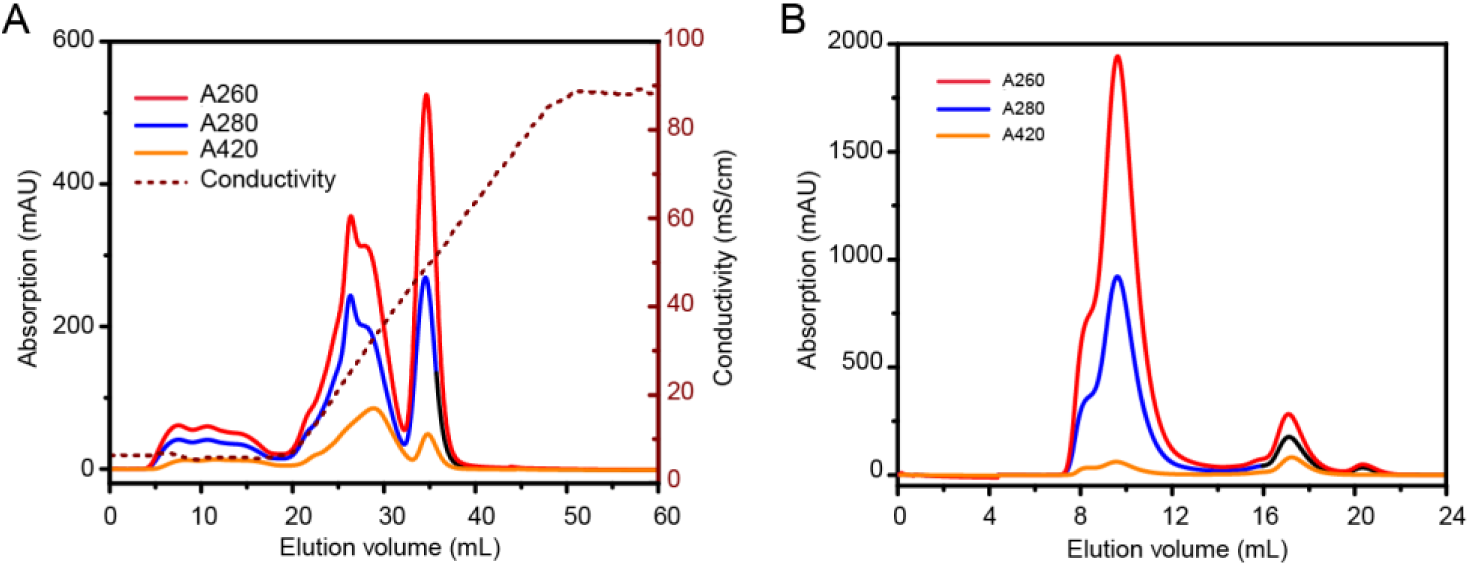
Purification of Nanogold-labeled RNA samples under native conditions. (*A*) The elution profile of Nanogold-labeled 3’SL RNA by anion exchange chromatography. (*B*) The elution profile of Nanogold-labeled DENV-mini RNA by size-exclusion chromatography. The elution curves corresponding to absorbance at specific wavelengths and conductivity are indicated in both A and B.

The successful conjugation of Nanogold to the TPT3^A^-modified RNAs can be verified by UV-vis absorbance at 420 nm. Generally, Nanogold exhibits prominent absorption at both 260 nm and 420 nm, natural RNA has strong absorption at 260 nm but very weak absorption at 420 nm. Thus, the absorption peak at 420 nm for the RNA conjugates indicate stable Nanogold coupling to RNAs (**Figs. 5A** and **5B**). The Nanogold coupling efficiency can be calculated from the absorption of the RNA conjugates at both 260 nm and 420 nm respectively using the respective molar extinction coefficients of RNAs and Nanogold (25). The efficiencies of the one site- and double-labeling range from 65% to 86% (***SI Appendix*, Table S6**).

### Distance Measurements by X-ray Scattering Interferometry

Measurements of molecular distances are key to dissect the structure, dynamics and functions of biomolecules (26). To demonstrate the applications of Nanogold-conjugated RNAs in large RNA structural biology, we measured the distance distributions between sites of A689 and U719 in the 3’SL and DENV-mini RNAs by XSI. SAXS experiments were performed for the sample quartets of 3’SL and DENV-mini RNA conjugates along with the free Nanogold. The scattering profiles normalized against I_0_ for the free Nanogold and the two orthogonal single-labeled, doubled-labeled Nanogold-RNA conjugates of 3’SL and DENV-mini are shown in **Figs. 6A-B**, respectively. In comparison with the scattering profiles of native 3’SL, the two single-labeled and the double-labeled Nanogold-RNA conjugates show systematic increase in intensity at elevated *q* range from 0.15 to 0.30 Å^-1^ (Fig. 6A), which is consistent with stable conjugation of the RNAs with one or two Nanogolds that scatter X-ray strongly. Similar pattern is observed for the DENV-mini RNAs (**Fig. 6B**). The overall structural parameters derived from scattering profiles including *R*_g_ and *D*_max_ are shown in ***SI Appendix*, Table S5**. The Rg and Dmax are slightly huge than native RNA, which can be a result of nanogold coupling, and also support that the nanogold labeling dose not substantially perturb RNA structure.

**Fig. 6.**
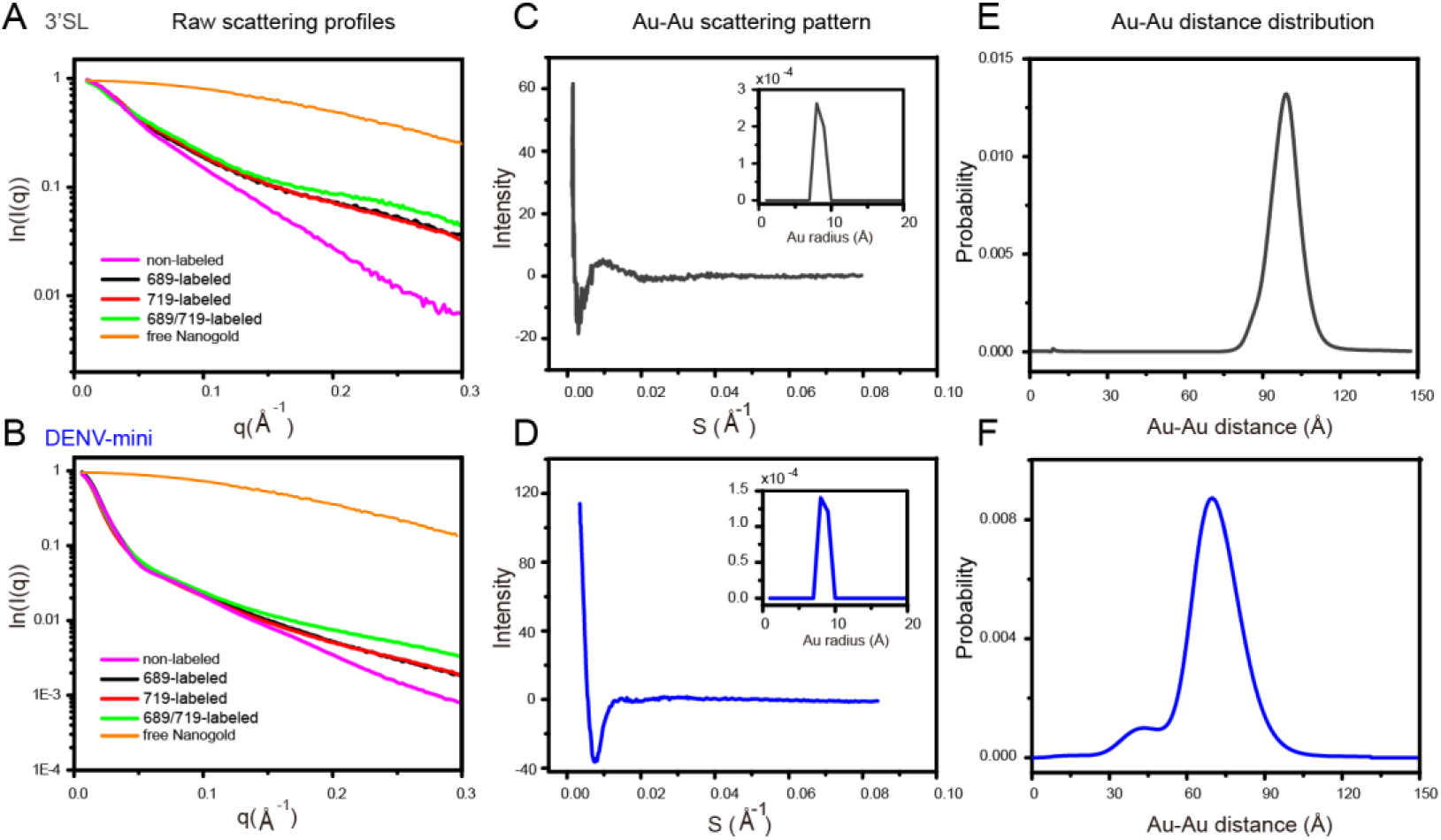
Measuring Nanogold-Nanogold distance distributions from Nanogold-conjugated 3’SL and DENV-mini RNA samples by X-ray scattering interferometry. (*A*-*B*) I_0_-normalizzed scattering profiles of free Nanogold (orange), 689-single labeled (black), 719-single labeled (red) and 689/719-double labeled (green) Nanogold-RNA conjugates, non-labeled native RNA (magenta) for 3’SL (A) and DENV-mini (B). (*C*-*D*) Scattering interference pattern of the two Nanogold labels in 3’SL (C) and DENV-mini (D). The insets are the radius distribution of the free Nanogold used in the experiment, showing a maximum probability at 7 Å. (*E*-*F*) Probability distribution of the center-to-center distance between the two Nanogold labeles in 3’SL (97.7 Å) (E) and DENV-mini (78.2 Å) (F), with variance of 52.7 Å^2^, 83.3 Å^2^, respectively. A shoulder peak with mean distance of 40 Å was observed for DENV-mini.

The full set of scattering profiles of the RNA quartets and the free Nanogold were combined to derive Nanogold-Nanogold scattering interference profiles (**Figs. 6C-D**) and the center-to-center distance distributions between the pairs of Nanogold (**Figs. 6E-F**), for the 3’SL and DENV-mini RNAs, respectively, by following the previously described protocols(20, 21). Both Nanogold-Nanogold scattering interference profiles show oscillating peaks, corresponding to major distance distributions with mean values of 97.7 Å, 78.2 Å and variance of 52.7 Å^2^, 83.3 Å^2^ for the 3’SL (**Figs. 6C** and **6E**) and DENV-mini (**Figs. 6D** and **6F**) RNA samples, respectively. The distance between N1 of U719 and N9 of A689 is measured as 83.3 Å in the atomic model of 3’SL derived from SAXS data (Fig. 1C). Given the radius of the Nanogold and the length of the linkers, the mean Nanogold-Nanogold distance of 97.7 Å in 3’SL RNA is reasonably consistent with the atomic model. The 20 Å decrease of the mean Nanogold-Nanogold distance between sites in DENV-mini RNA supports conformational changes in 3’SL upon genome circularization mediated by long-range RNA-RNA interactions between the 5’ and 3’ ends. The larger variance of the distance distribution in DENV-mini RNA suggests that 3’SL element becomes more flexible and may adopt an ensemble of conformations upon genome circularization, which is further evidenced by the small shoulder peak with mean distance of 40 Å (**Fig. 6F**).

## Discussion

In this work, we present an easy, efficient and generally applicable labeling strategy for site-specific covalent conjugation of large RNAs with gold nanoparticle empowered by expanded genetic alphabet transcription. By site-specific labeling of a 97-nt 3’SL and a 719-nt DENV-mini RNAs with one or a pair of Nanogold, we measured the inter-Nanogold distance distributions between the labeling sites with XSI and validate the labeling strategy. For the first time, we provide experimental evidence for tertiary conformational changes in such large RNAs caused by long-range RNA-RNA interactions, thus supporting the hypothetical flavivirus genome circularization model. This labeling strategy overcomes the size constraints in conventional RNA labeling methods. In principle, it can be applied to RNAs with sizes from tens up to thousands of nucleotides due to the high processivity of T7 RNA polymerase. The design of a far upstream forward primer enables easy and efficient native purification of the RNA transcripts from DNA templates and excess nucleotides. Since all the reactions are performed in near physiological conditions, it avoids the denaturing steps in other labeling strategies and help preserve the native conformations of large RNAs for subsequent analysis.

The presented strategy is expected to promote wide applications of gold-nanoparticle-RNA conjugates in large RNA structural biology. To our knowledge, the 97-nt 3’SL and the 719-nt DENV-mini RNAs are among the largest ones that have been site-specifically conjugated with gold nanoparticles and studied by XSI so far. As XSI can measure much broader distance distributions (ranging from 50 up to 400 Å) than other molecular rulers, such as pulsed electron paramagnetic resonance spectroscopy (EPR) and Förster resonance energy transfer (FRET) that can provide distance information from 20-80 Å (18), XSI can play important roles in structural study of large RNAs and RNA complexes, such as the long non-coding RNAs. For example, the XSI-derived distance distributions can be used to aid in computational 3D modeling of large RNAs (16). Furthermore, we envision that gold nanoparticle-RNA conjugates can be utilized in studying the structure and dynamics of RNAs by cryo-EM microscopy, an EM-based technique called individual-particle electron tomography, or high-resolution AFM (9, 48). Very recently, the structure and motion of the long noncoding RNA HOTAIR (2,158 nts) have been visualized by AFM, but better anatomy assignments could be achieved with AFM imaging of site-specifically gold nanoparticle-labeled RNA (RNA 2020, doi: 10.1261/rna.074633.120.). The presented labeling strategy could work for such a large RNA.

The presented strategy could also have wide applications in RNA nanotechnology (4). Recently, several fluorogenic or cell-specific RNA aptamers have been developed for live cell applications (4). Bioconjugation of RNA aptamers with nanoparticles could combine unique and orthogonal strengths of the specific interactions of RNA nanostructures and novel material properties of nanoparticles and open up new options for a wide range of applications (10). For example, the utility of SAXS, AFM for analysis of nanoparticle-conjugated RNA nanostructures could also be established (13). RNA aptamer-nanoparticle conjugates could be powerful diagnostic and therapeutic tools (49).

Last but not least, the presented strategy can be used for site-specific conjugation of RNAs with other metal nanoparticles including silver, titanium oxide, zinc, ion nanoparticles for diverse biomedical applications. The expanded genetic alphabet transcription also allows for flexible chemistry for site-specific covalent conjugation. For example, alkyne- or azide-derivatized UBP (e.g. TPT3) can be synthesized and incorporated into RNAs by expanded genetic alphabet transcription for subsequent site-specific conjugation with commercial Click Nanogold probes (http://www.nanoprobes.com/) *via* click chemistry (31, 32).

## Materials and Methods

### Material

The deoxyribonucleotide phosphoramidites (dNaM-CEP/dTPT3-CEP), triphosphorylated deoxy-aribonucleotides (dTPT3TP/dNaMTP) and ribonucleotides (rTPT3TP/rNaMTP) were synthesized according to the literature procedures (31, 33, 34). 2× Golden mix of Taq DNA polymerase (with buffer and natural dNTPs mix included) was purchased from TSINGKE Biological Technology Co., Ltd. (Beijing, China). The T7 RNA polymerase was homemade. The 1.4 nm Mono-Sulfo-NHS-Nanogold, with the molar extinction coefficient at 420 nm being 155,000 M^-1^cm^-1^, was purchased from Nanoprobes (http://www.nanoprobes.com/). Plasmids encoding the 3’SL or the DENV-mini RNAs of DENV2 with an upstream T7 promoter were total-gene synthesized and sequenced by Wuxi Qinglan Biotechnology Inc, Wuxi, China. All ssDNA primers containing natural or unnatural nucleotides were synthesized and purified with OPC purification by TSINGKE. The DNA sequences of the plasmids and the natural and unnatural primers in this work could be found in ***SI Appendix*, Tables S1, S2**, respectively.

### Preparation of Native or UBP-modified DNA/RNAs

With the synthesized plasmids as templates and directed by the respective pairs of primers (***SI Appendix*, Table S2**), the dsDNA templates for the respective RNA constructs were generated by PCR reactions (***SI Appendix*, Table S3**). Prior to large scale RNA sample preparation, the optimal conditions for *in vitro* transcription were screened against Mg^2+^ and DNA template concentrations. The same NaM-containing dsDNA templates were used for incorporation of rTPT3 or rTPT3^A^ into 3’SL and DENV-mini RNAs by *in vitro* transcription (***SI Appendix*, Table S4**). The transcription supernatants were directly applied to a HiLoad 16/600 Superdex 75 (for 3’SL) or 200 (for DENV-mini) gel filtration columns and the RNAs were purified by size exclusion chromatography (SEC). The SEC buffer contains 20 mM HEPES pH 7.5, 100 mM KCl, 5 mM Mg^2+^. RNA fractions were collected and concentrated with the Amicon Ultra centrifugal filter devices (Sigma-Aldrich) and stored at −80 °C for further use. The concentrations of native or Nanogold-labeled RNA samples were determined by UV-vis absorbance at 260 nm or 420 nm, respectively, on a NanoDrop 2000 (Thermo Scientific). The molar extinction coefficients of RNAs at 260 nm were calculated from sequences with the OligoAnalyzer Tool (https://sg.idtdna.com/pages/tools/oligoanalyzer).

### Site-specific Nanogold Labeling of RNAs

The purified single-site or double TPT3^A^-modified 3’SL and DENV-mini RNAs were buffer-exchanged into 0.1 M NaHCO_3_, pH 8.0 and subjected to Nanogold labeling. 1.4 nm Nanogold was dissolved with DEPC H_2_O following the product instruction manuals (http://www.nanoprobes.com/). The coupling reactions were initiated by mixing 8 nM single- or double-site TPT3^A^-modified RNA samples with 16 nM or 32 nM 1.4 nm Nanogold, respectively, then incubated at 25°C overnight. To increase the solubility and coupling efficiency of Nanogold, 0.05% (v/v) Triton X-100 was added in the reaction buffer. Nanogold-RNA conjugates are further purified after the reactions are done. For 3’SLs, the reaction mixtures were first injected into Hitrap Q column (GE Healthcare) pre-equilibrated with buffer A (20 mM HEPES, 20 mM KCl, 5 mM Mg^2+^, 3% glycerol and 1 mM TCEP, pH 7.5), and then eluted with a salt gradient to 100% buffer B (20 mM HEPES, 500 mM KCl, 5 mM Mg^2+^, 3% glycerol and 1 mM TCEP, pH 7.5). Free Nanogold and Nanogold-labeled 3’SL RNAs were eluted at about 20% and 50% Buffer B, respectively. For Nanogold-labeled DENV-mini RNAs, the free Nanogold can be separated directly by passing the reaction mixture through a Superdex 200 gel filtration column.

### Small Angle X-ray Scattering

All the natural and unnatural RNA samples, as well as the Nanogold-labeled RNAs and free Nanogold, were exchanged into final SAXS buffer (20 mM HEPES pH 7.5, 100 mM KCl, 5 mM Mg^2+^, 0.5 mM TCEP, 3% (v/v) glycerol) using SEC columns or Amicon Ultra centrifugal filter devices (Millipore). All natural and unnatural RNA samples without labelling were diluted to final concentrations of 0.25-1 mg/ml, all Nanogold-labeled RNA solutions were concentrated and adjusted to 30 µM, and the free Nanogold sample was concentrated to 100 μM. The parameters for data collection and software employed for data analysis are summarize in ***SI Appendix***, Table S7.

The data collection and processing procedures are similar as described before (39). Briefly, SAXS measurements were carried out at room temperature at the beamline 12 ID-B of the Advanced Photon Source, Argonne National Laboratory. The setups were adjusted to achieve scattering q values of 0.005 < q < 0.89 Å-1 (12ID-B), where q = (4π/λ) sinθ, and 2θ is the scattering angle. Thirty two-dimensional images were recorded for each buffer or sample solution using a flow cell, with the exposure time of 1 s for natural RNA samples, or of 0.1 s for Nanogold-labeled RNA samples so as to minimize radiation damage and obtain good signal-to-noise ratio. The 2D images were reduced to one-dimensional scattering profiles by using Matlab onsite. The scattering profiles, the forward scattering intensity I(0) and the radius of gyration (*R*_g_), the pair distance distribution function (PDDF) p(r) as well as the maximum dimension, *D*_max_ of the RNAs were calculated using same procedures as described before (39). The Volume-of-correlation (Vc) were calculated by using the program Scatter, and the molecular weights of natural RNAs were calculated on a relative scale using the Rg/Vc power law developed by Rambo et al (47), independent of RNA concentrations and with minimal user bias.

### XSI Data Processing

The scattering profiles of the RNA quartets for 3’SL and DENV-mini RNAs and the free Nanogold were used to calculate the Nanogold-Nanogold scattering interference profiles, from which the Nanogold-Nanogold distance distributions were inferred with the previously described protocols (20, 21) and the Matlab scripts in AuSAXSGUI (https://github.com/thomas836/AuSAXSGUI) shared by Jan Lipfert group. Briefly, after standard SAXS data processing, the radius distribution of Nanogold is firstly determined from the free Nanogold scattering profile. The Nanogold-Nanogold scattering interference profiles, I_Au-Au(S)_, is then calculated as the sum of the concentration-normalized scattering signals of the double-labeled samples and non-labeled sample minus the signals of the two single-labeled samples. Systems with two Nanogold separated at different distances will give different interference profiles. The experimentally obtained I_Au-Au(S)_ is then decomposed into contributions from 200 uniformly spaced Nanogold-Nanogold distances of 1 to 200 Å to generate the Nanogold-Nanogold distance probability distribution using a procedure that maximize the entropy.

## Acknowledgments

This work was supported by grants from the National Natural Science Foundation of China (No. U1832215), the Beijing Advanced Innovation Center for Structural Biology, the Tsinghua-Peking Joint Center for Life Sciences to X.F.. We thank Dr. Xiaobing Zuo at the beamline 12-ID-B, Advanced Photon Source, Argonne National Laboratory, USA for assistance during data collection.

## References

1. J. M. Engreitz, N. Ollikainen, M. Guttman, Long non-coding RNAs: spatial amplifiers that control nuclear structure and gene expression. Nat Rev Mol Cell Biol 17, 756–770 (2016).

2. V. Bernat, M. D. Disney, RNA Structures as Mediators of Neurological Diseases and as Drug Targets. Neuron 87, 28–46 (2015).

3. H. Ohno, S. Akamine, H. Saito, RNA nanostructures and scaffolds for biotechnology applications. Curr Opin Biotechnol 58, 53–61 (2019).

4. D. Jasinski, F. Haque, D. W. Binzel, P. Guo, Advancement of the Emerging Field of RNA Nanotechnology. ACS Nano 11, 1142–1164 (2017).

5. E. Paredes, M. Evans, S. R. Das, RNA labeling, conjugation and ligation. Methods 54, 251–259 (2011).

6. J. T. George, S. G. Srivatsan, Posttranscriptional chemical labeling of RNA by using bioorthogonal chemistry. Methods 120, 28–38 (2017).

7. P. M. Tiwari, K. Vig, V. A. Dennis, S. R. Singh, Functionalized Gold Nanoparticles and Their Biomedical Applications. Nanomaterials (Basel) 1, 31–63 (2011).

8. D. A. Giljohann et al., Gold nanoparticles for biology and medicine. Angew Chem Int Ed Engl 49, 3280–3294 (2010).

9. R. D. Powell, J. F. Hainfeld, Preparation and high-resolution microscopy of gold cluster labeled nucleic acid conjugates and nanodevices. Micron 42, 163–174 (2011).

10. I. Tessmer, P. Kaur, J. Lin, H. Wang, Investigating bioconjugation by atomic force microscopy. J Nanobiotechnology 11, 25 (2013).

11. C. A. Brosey, J. A. Tainer, Evolving SAXS versatility: solution X-ray scattering for macromolecular architecture, functional landscapes, and integrative structural biology. Curr Opin Struct Biol 58, 197–213 (2019).

12. X. Fang, J. R. Stagno, Y. R. Bhandari, X. Zuo, Y. X. Wang, Small-angle X-ray scattering: a bridge between RNA secondary structures and three-dimensional topological structures. Curr Opin Struct Biol 30, 147–160 (2015).

13. M. A. B. Baker et al., Dimensions and Global Twist of Single-Layer DNA Origami Measured by Small-Angle X-ray Scattering. ACS Nano 12, 5791–5799 (2018).

14. L. K. Bruetzel et al., Conformational Changes and Flexibility of DNA Devices Observed by Small-Angle X-ray Scattering. Nano Lett 16, 4871–4879 (2016).

15. S. Fischer et al., Shape and Interhelical Spacing of DNA Origami Nanostructures Studied by Small-Angle X-ray Scattering. Nano Lett 16, 4282–4287 (2016).

16. A. Ponce-Salvatierra et al., Computational modeling of RNA 3D structure based on experimental data. Biosci Rep 39 (2019).

17. R. S. Mathew-Fenn, R. Das, P. A. Harbury, Remeasuring the double helix. Science 322, 446–449 (2008).

18. R. S. Mathew-Fenn, R. Das, J. A. Silverman, P. A. Walker, P. A. Harbury, A molecular ruler for measuring quantitative distance distributions. PLoS One 3, e3229 (2008).

19. T. Zettl et al., Gold nanocrystal labels provide a sequence-to-3D structure map in SAXS reconstructions. Sci Adv 4, eaar4418 (2018).

20. X. Shi, S. Bonilla, D. Herschlag, P. Harbury, Quantifying Nucleic Acid Ensembles with X-ray Scattering Interferometry. Methods Enzymol 558, 75–97 (2015).

21. T. Zettl et al., Recording and Analyzing Nucleic Acid Distance Distributions with X-Ray Scattering Interferometry (XSI). Curr Protoc Nucleic Acid Chem 73, e54 (2018).

22. G. L. Hura et al., DNA conformations in mismatch repair probed in solution by X-ray scattering from gold nanocrystals. Proc Natl Acad Sci U S A 110, 17308–17313 (2013).

23. X. Shi, L. Huang, D. M. Lilley, P. B. Harbury, D. Herschlag, The solution structural ensembles of RNA kink-turn motifs and their protein complexes. Nat Chem Biol 12, 146–152 (2016).

24. X. Shi, P. Walker, P. B. Harbury, D. Herschlag, Determination of the conformational ensemble of the TAR RNA by X-ray scattering interferometry. Nucleic Acids Res 45, e64 (2017).

25. C. J. Ackerson, R. D. Powell, J. F. Hainfeld, Site-specific biomolecule labeling with gold clusters. Methods Enzymol 481, 195–230 (2010).

26. T. Zettl et al., Absolute Intramolecular Distance Measurements with Angstrom-Resolution Using Anomalous Small-Angle X-ray Scattering. Nano Lett 16, 5353–5357 (2016).

27. I. Lebars et al., A fully enzymatic method for site-directed spin labeling of long RNA. Nucleic Acids Res 42, e117 (2014).

28. E. S. Babaylova et al., Complementary-addressed site-directed spin labeling of long natural RNAs. Nucleic Acids Res 44, 7935–7943 (2016).

29. M. Zhao et al., Site-specific dual-color labeling of long RNAs for single-molecule spectroscopy. Nucleic Acids Res 46, e13 (2018).

30. D. A. Malyshev, F. E. Romesberg, The expanded genetic alphabet. Angew Chem Int Ed Engl 54, 11930–11944 (2015).

31. Y. J. Seo, D. A. Malyshev, T. Lavergne, P. Ordoukhanian, F. E. Romesberg, Site-specific labeling of DNA and RNA using an efficiently replicated and transcribed class of unnatural base pairs. J Am Chem Soc 133, 19878–19888 (2011).

32. T. Someya, A. Ando, M. Kimoto, I. Hirao, Site-specific labeling of RNA by combining genetic alphabet expansion transcription and copper-free click chemistry. Nucleic Acids Res 43, 6665–6676 (2015).

33. L. Li et al., Natural-like replication of an unnatural base pair for the expansion of the genetic alphabet and biotechnology applications. J Am Chem Soc 136, 826–829 (2014).

34. Y. J. Seo, G. T. Hwang, P. Ordoukhanian, F. E. Romesberg, Optimization of an unnatural base pair toward natural-like replication. J Am Chem Soc 131, 3246–3252 (2009).

35. L. G. Gebhard, C. V. Filomatori, A. V. Gamarnik, Functional RNA elements in the dengue virus genome. Viruses 3, 1739–1756 (2011).

36. N. G. Iglesias, A. V. Gamarnik, Dynamic RNA structures in the dengue virus genome. RNA Biol 8, 249–257 (2011).

37. B. L. Nicholson, K. A. White, Functional long-range RNA-RNA interactions in positive-strand RNA viruses. Nat Rev Microbiol 12, 493–504 (2014).

38. M. A. Brinton, M. Basu, Functions of the 3’ and 5’ genome RNA regions of members of the genus Flavivirus. Virus Res 206, 108–119 (2015).

39. Y. Zhang et al., Long non-coding subgenomic flavivirus RNAs have extended 3D structures and are flexible in solution. EMBO Rep 10.15252/embr.201847016, e47016 (2019).

40. S. You, R. Padmanabhan, A novel in vitro replication system for Dengue virus. Initiation of RNA synthesis at the 3’-end of exogenous viral RNA templates requires 5’- and 3’-terminal complementary sequence motifs of the viral RNA. J Biol Chem 274, 33714–33722 (1999).

41. J. Sztuba-Solinska et al., Structural complexity of Dengue virus untranslated regions: cis-acting RNA motifs and pseudoknot interactions modulating functionality of the viral genome. Nucleic Acids Res 41, 5075–5089 (2013).

42. Y. J. Seo, S. Matsuda, F. E. Romesberg, Transcription of an expanded genetic alphabet. J Am Chem Soc 131, 5046–5047 (2009).

43. T. Lavergne et al., FRET Characterization of Complex Conformational Changes in a Large 16S Ribosomal RNA Fragment Site-Specifically Labeled Using Unnatural Base Pairs. ACS Chem Biol 11, 1347–1353 (2016).

44. S. N. Ho, H. D. Hunt, R. M. Horton, J. K. Pullen, L. R. Pease, Site-directed mutagenesis by overlap extension using the polymerase chain reaction. Gene 77, 51–59 (1989).

45. Y. Zhang et al., A semi-synthetic organism that stores and retrieves increased genetic information. Nature 551, 644–647 (2017).

46. A. X. Zhou, K. Sheng, A. W. Feldman, F. E. Romesberg, Progress toward Eukaryotic Semisynthetic Organisms: Translation of Unnatural Codons. J Am Chem Soc 141, 20166–20170 (2019).

47. R. P. Rambo, J. A. Tainer, Accurate assessment of mass, models and resolution by small-angle scattering. Nature 496, 477–481 (2013).

48. L. Zhang et al., Three-dimensional structural dynamics and fluctuations of DNA-nanogold conjugates by individual-particle electron tomography. Nat Commun 7, 11083 (2016).

49. H. Jo, C. Ban, Aptamer-nanoparticle complexes as powerful diagnostic and therapeutic tools. Exp Mol Med 48, e230 (2016).

