## Supporting information for "Site-specific covalent labeling of large RNAs with nanoparticles empowered by expanded genetic alphabet transcription"

### Site-specific Nanogold labeling of large RNAs with an unnatural base pair system under non-denaturing conditions for XSI distance measurements

Yan Wang[a], Yaoyi Chen[a, b], Yanping Hu[a], Xianyang Fang[a, *]

[a]Beijing Advanced Innovation Center for Structural Biology, School of Life Sciences, Tsinghua University, Beijing 100084, China

[b]Present address: Department of Mathematics and Computer Science, Freie Universität Berlin, Berlin 14195, Germany

#### This PDF file includes

- Supplementary Figures S1-S3
- Supplementary Tables S1-S7
- Synthetic procedures and characterizations of $rTPT3^{A}$

#### Supplementary Figures

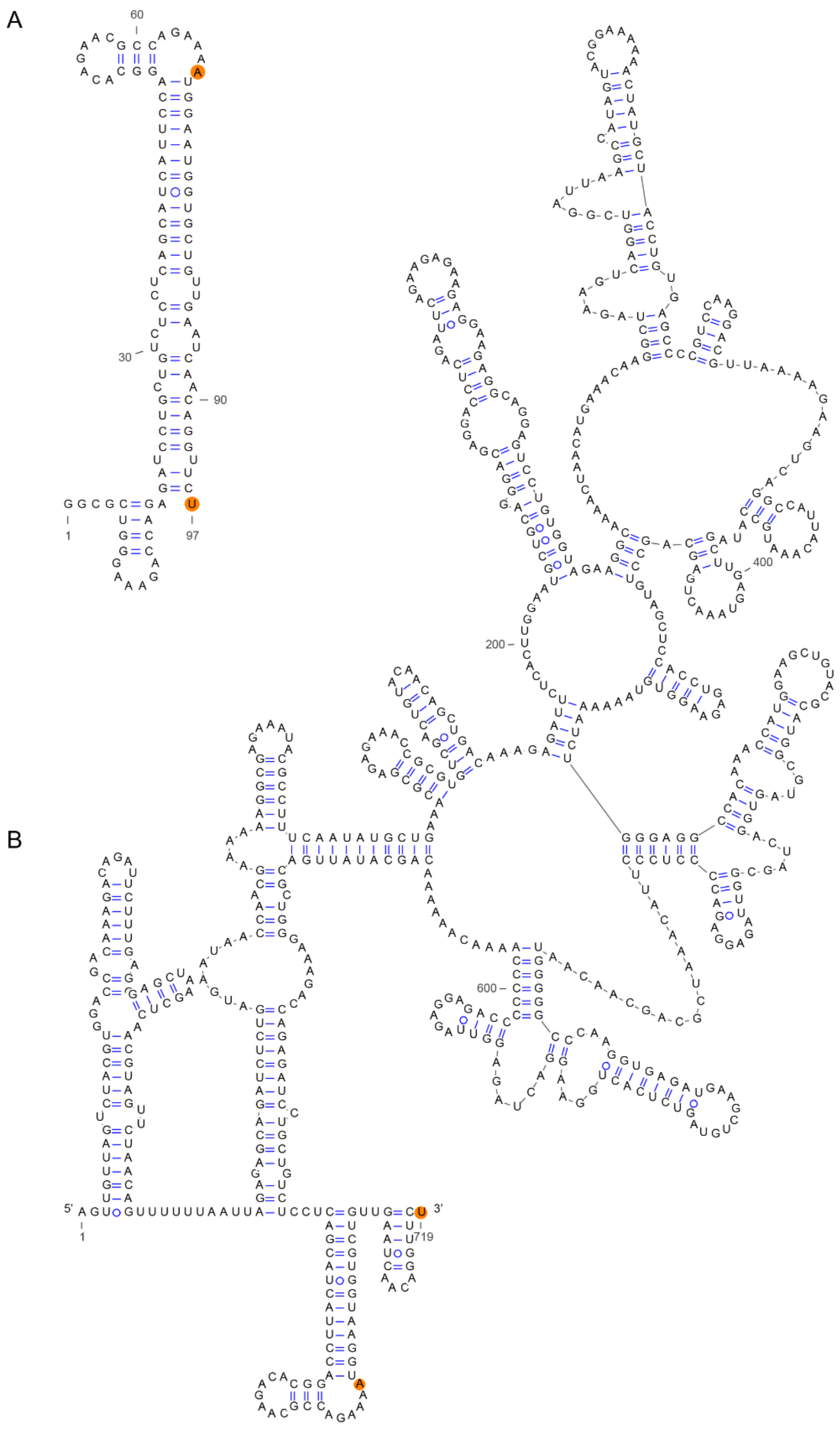


**Figure S1**. (**A-B**) Secondary structures of the 3'SL (**A**) and DENV-mini (**B**) RNAs used in this study. The secondary structure of DENV-mini is the same as in literature[1].

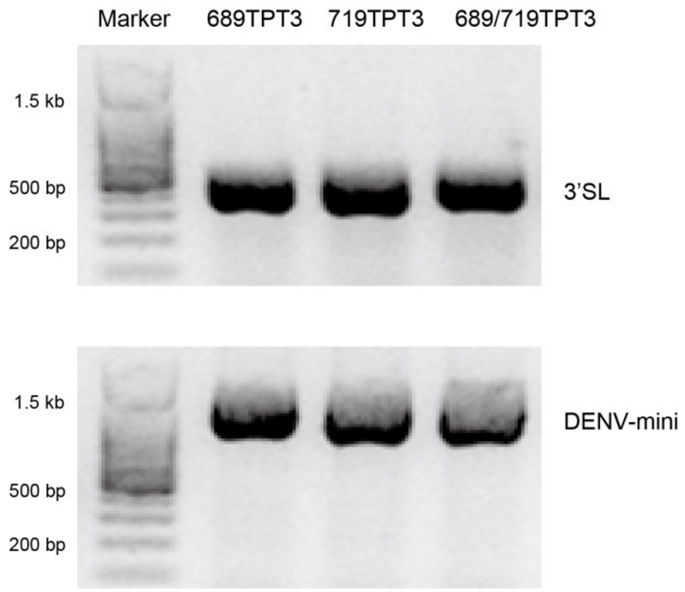


**Figure S2**. 1% Agrose gel analysis of the 689-single modified, 719-single modified, and 689/719-double modified PCR products coding for 3'SL (504 bp, top) and DENV-mini (1128 bp, down).

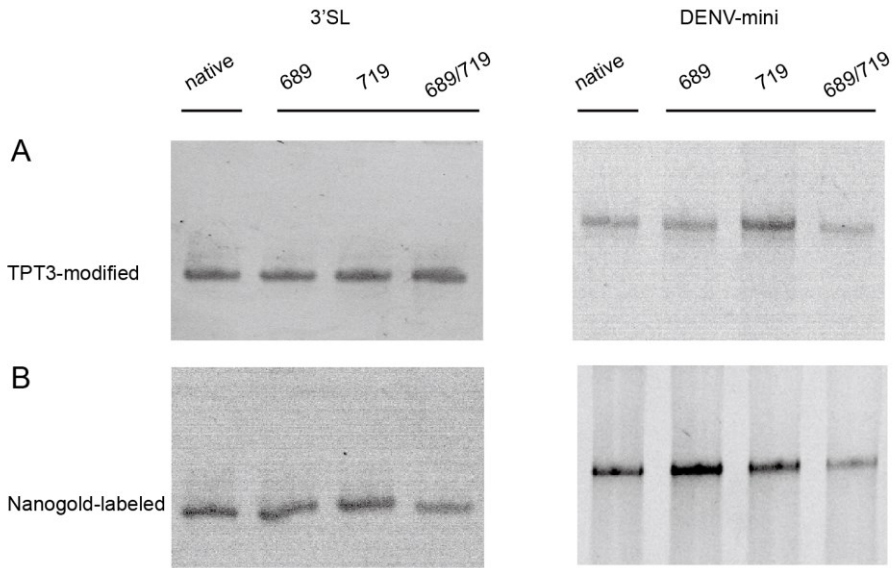


**Figure S3**. (**A-B**) 10% (left, 3’SL) and 5% (right, DENV-mini) PAGE analysis of purified TPT3-modified RNAs (**A**) and Nanogold-labeled RNAs (**B**). The respective lanes are indicated as WT (for non-labeled native RNA), 689 (for modified or labeled at site 689), 719 (for modified or labeled at site 719), 689/719 (for modified or labeled at both sites of 689 and 719).

### Supplementary Tables

**Table S1. The DNA sequences of the total gene synthesized plasmids coding for full-length 3'SL and DENV-mini RNAs from DENV2.**

| Plasmids | Primary sequence 5'-3' |
|---|---|
| 3'SL | AAAAATAAACAAATAGGGGTTCCGCGCACATTTCCCCGAAAAGTGCCACCTGA<br>CGTCTAAGAAACCATTATTATCATGACATTAACCTATAAAAATAGGCGTATCACG<br>AGGCCCTTTCGTTGTAAAACGACGGCCAGTCCGTCTCTATCCGGTCTCGATCCG<br>CAGTCTCTTGGCACAGGTAATACGACTCACTATAGGCGCTGGGAAAGACCAGA<br>GATCCTGCTGTCTCCTCAGCATCATTCCAGGCACAGAACGCCAGAAAATGGAAT<br>GGTGCTGTTGAATCAACAGGTTCTACCGTGCGAGTTACTGCCAACCGAGACCC<br>AACCGAGACGGGTCATAGCTGTTTCCAGTGTGCCGCTTCCTCGCTCACTGACTC<br>GCTGCGCTCGGTCGTTCGGCTGCGGCGAGCGGTATCAGCTCACTCAAAGGCGG<br>TAATACGGTTACCCACAGAATCAGGGGATAACGCAGGAAAGAACATGTGAGCA<br>AAAGGCCAGCAAAAGGCCAGGAACCGTAAAAAGGCCGCGTTGCTGGCGTTTT<br>TCCATAGGCTCCGCCCCCCTGACGAGCATCACAAAAATCGACGCTCAAGTCAG<br>AGGTGGCGAAACCCGACAGGACTATAAAGATACCAGGCGTTTCCCCCTGGAAG<br>CTCCCTCGTGCGCTCTCCTGTTCCGACCCTGCCGCTTACCGGATACCTGTCCGCC<br>TTTCTCCCTTCGGGAAGCGTGGCGCTTTCTCAATGCTCACGCTGTAGGTATCTCA<br>GTTCGGTGTAGGTCGTTCGCTCCAAGCTGGGCTGTGTGCACGAACCCCCCGTTC<br>AGCCCGACCGCTGCGCCTTATCCGGTAACTATCGTCTTGAGTCCAACCCGGTAA<br>GACACGACTTATCGCCACTGGCAGCAGCCACTGGTAACAGGATTAGCAGAGCG<br>AGGTATGTAGGCGGTGCTACAGAGTTCTTGAAGTGGTGGCCTAACTACGGCTAC<br>ACTAGAAGGACAGTATTTGGTATCTGCGCTCTGCTGAAGCCAGTTACCTTCGGA<br>AAAAGAGTTGGTAGCTCTTGATCCGGCAAACAAACCACCGCTGGTAGCGGTGG<br>TTTTTTTGTTTGCAAGCAGCAGATTACGCGCAGAAAAAAAGGATCTCAAGAAG<br>ATCCTTTGATCTTTTCTACGGGGTCTGACGCTCAGTGGAACGAAAACTCACGTT<br>AAGGGATTTTGGTCATGAGATTATCAAAAAGGATCTTCACCTAGATCCTTTTAAA<br>TTAAAAATGAAGTTTTAAATCAATCTAAAGTATATATGAGTAAACTTGGTCTGAC<br>AGTTACCAATGCTTAATCAGTGAGGCACCTATCTCAGCGATCTGTCTATTTCGTT<br>CATCCATAGTTGCCTGACTCCCCGTCGTGTAGATAACTACGATACGGGAGGGCT<br>TACCATCTGGCCCCAGTGCTGCAATAATACCGCGGGACCCACGCTCACCGGCTC<br>CAGATTTATCAGCAATAAACCAGCCAGCCGGAAGGGCCGAGCGCAGAAGTGGT<br>CCTGCAACTTTATCCGCCTCCATCCAGTCTATTAATTGTTGCCGGGAAGCTAGAG<br>TAAGTAGTTCGCCAGTTAATAGTTTGCGCAACGTTGTTGCCATCGCTACAGGCAT<br>CGTGGTGTCACGCTCGTCGTTTGGTATGGCTTCATTCAGCTCCGGTTCCCAACG<br>ATCAAGGCGAGTTACATGATCCCCCATGTTGTGCAAAAAAGCGGTTAGCTCCTT<br>CGGTCCTCCGATCGTTGTCAGAAGTAAGTTGGCCGCCGTGTTATCACTCATGGT<br>TATGGCAGCACTACATAATTCTCTTACTGTCATGCCATCCGTAAGATGCTTTTCTG<br>TGACTGGTGAGTACTCAACCAAGTCATTCTGAGAATAGTGTATGCGGCGACCGA<br>GTTGCTCTTGCCCGGCGTCAATACGGGATAATACCGCGCCACATAGCAGAACTT<br>TAAAAGTGCTCATCATTGGAAAACGTTCTTCGGGGCGAAAACTCTCAAGGATCT<br>TACCGCTGTTGAGATCCAGTTCGATGTAACCCACTCGTGCACCCAACTGATCTT<br>CAGCATCTTTTACTTTCACCAGCGTTTCTGGGTGAGCAAAAACAGGAAGGCAA<br>AATGCCGCAAAAAAGGGAATAAGGGCGACACGGAAATGTTGAATACTCATACT<br>CTTCCTTTTTCAATATTATTGAAGCATTTATCAGGGTTATTGTCTCATGAGCGGAT<br>ACATATTTGAATGTATTTAG |
| DENV-mini | AAAAATAAACAAATAGGGGTTCCGCGCACATTTCCCCGAAAAGTGCCACCTGA<br>CGTCTAAGAAACCATTATTATCATGACATTAACCTATAAAAATAGGCGTATCACG<br>AGGCCCTTTCGTTGTAAAACGACGGCCAGTCGAACCACGCAATGCGTCTCGAT<br>CCGCAGTGTCTTGCGTCTCTTAATACGACTCACTATAAGTTGTTAGTCTACGTGG<br>ACCGACAAAGACAGATTCTTTGAGGGAGCTAAGCTCAACGTAGTTCTAACAGT<br>TTTTTAATTAGAGAGCAGATCTCTGATGAATAACCAACGAAAAAAGGCGAGAA<br>ATACGCCTTTCAATATGCTGAAACGCGAGAGAAACCGCGTGTCGACTGTACAAC<br>AGCTGACAAAGAGATTCTCACTTGGAATGCTGCAGGGACGAGGACCTCAGATT<br>CAGAAGAGAAGAGGAAGAGGCAGGAGTCCTGTGGTAGAAGGCAAAACTAACA<br>TGAAACAAGGCTAGAAGTCAGGTCGGATTAAGCCATAGTACGGAAAAAACTAT<br>GCTACCTGTGAGCCCCGTCCAAGGACGTTAAAAGAAGTCAGGCCATTACAAAT |

| | |
|---|---|
| | GCCATAGCTTGAGTAAACTGAGCAGCCTGTAGCTCCACCTGAGAAGGTGTAAA<br>AAATCTGGGAGGCCACAAACCATGGAAGCTGTACGCATGGCGTAGTGGACTAG<br>CGGTTAGAGGAGACCCCTCCCTTACAAATCGCAGCAACAATGGGGGCCCAAGG<br>TGAGATGAAGCTGTAGTCTCACTGGAAGGACTAGAGGTTAGAGGAGACCCCCC<br>CAAAACAAAAAACAGCATATTGACGCTGGGAAAGACCAGAGATCCTGCTGTCT<br>CCTCAGCATCATTCCAGGCACAGAACGCCAGAAAATGGAATGGTGCTGTTGAA<br>TCAACAGGTTCTAGAGACGGAGTCACTGCCAACCGAGACGGTCATAGCTGTTT<br>CCTGTGTGCCGCTTCCTCGCTCACTGACTCGCTGCGCTCGGTCGTTCGGCTGCG<br>GCGAGCGGTATCAGCTCACTCAAAGGCGGTAATACGGTTACCCACAGAATCAG<br>GGGATAACGCAGGAAAGAACATGTGAGCAAAAGGCCAGCAAAAGGCCAGGAA<br>CCGTAAAAAGGCCGCGTTGCTGGCGTTTTTCCATAGGCTCCGCCCCCCTGACGA<br>GCATCACAAAAATCGACGCTCAAGTCAGAGGTGGCGAAACCCGACAGGACTAT<br>AAAGATACCAGGCGTTTCCCCCTGGAAGCTCCCTCGTGCGCTCTCCTGTTCCGA<br>CCCTGCCGCTTACCGGATACCTGTCCGCCTTTCTCCCTTCGGGAAGCGTGGCGC<br>TTTCTCATAGCTCACGCTGTAGGTATCTCAGTTCGGTGTAGGTCGTTCGCTCCAA<br>GCTGGGCTGTGTGCACGAACCCCCCGTTCAGCCCGACCGCTGCGCCTTATCCG<br>GTAACTATCGTCTTGAGTCCAACCCGGTAAGACACGACTTATCGCCACTGGCAG<br>CAGCCACTGGTAACAGGATTAGCAGAGCGAGGTATGTAGGCGGTGCTACAGAG<br>TTCTTGAAGTGGTGGCCTAACTACGGCTACACTAGAAGGACAGTATTTGGTATC<br>TGCGCTCTGCTGAAGCCAGTTACCTTCGGAAAAAGAGTTGGTAGCTCTTGATCC<br>GGCAAACAAACCACCGCTGGTAGCGGTGGTTTTTTTGTTTGCAAGCAGCAGAT<br>TACGCGCAGAAAAAAAGGATCTCAAGAAGATCCTTTGATCTTTTCTACGGGGTC<br>TGACGCTCAGTGGAACGAAAACTCACGTTAAGGGATTTTGGTCATGAGATTATC<br>AAAAAGGATCTTCACCTAGATCCTTTTAAATTAAAAATGAAGTTTTAAATCAATC<br>TAAAGTATATATGAGTAAACTTGGTCTGACAGTTACCAATGCTTAATCAGTGAGG<br>CACCTATCTCAGCGATCTGTCTATTTCGTTCATCCATAGTTGCCTGACTCCCCGTC<br>GTGTAGATAACTACGATACGGGAGGGCTTACCATCTGGCCCCAGTGCTGCAATA<br>ATACCGCGGGACCCACGCTCACCGGCTCCAGATTTATCAGCAATAAACCAGCCA<br>GCCGGAAGGGCCGAGCGCAGAAGTGGTCCTGCAACTTTATCCGCCTCCATCCA<br>GTCTATTAATTGTTGCCGGGAAGCTAGAGTAAGTAGTTCGCCAGTTAATAGTTTG<br>CGCAACGTTGTTGCCATCGCTACAGGCATCGTGGTATCACGCTCGTCGTTTGGT<br>ATGGCTTCATTCAGCTCCGGTTCCCAACGATCAAGGCGAGTTACATGATCCCCC<br>ATGTTGCGCAAAAAAGCGGTTAGCTCCTTCGGTCCTCCGATCGTTGTCAGAAGT<br>AAGTTGGCCGCCGTGTTATCACTCATGGTTATGGCAGCACTACATAATTCTCTTA<br>CTGTCATGCCATCCGTAAGATGCTTTTCTGTGACTGGTGAGTACTCAACCAAGT<br>CATTCTGAGAATAGTGTATGCGGCGACCGAGTTGCTCTTGCCCGGCGTCAATAC<br>GGGATAATACCGCGCCACATAGCAGAACTTTAAAAGTGCTCATCATTGGAAAAC<br>GTTCTTCGGGGCGAAAACTCTCAAGGATCTTACCGCTGTTGAGATCCAGTTCGA<br>TGTAACCCACTCGTGCACCCAACTGATCTTCAGCATCTTTTACTTTCACCAGCGT<br>TTCTGGGTGAGCAAAAACAGGAAGGCAAAATGCCGCAAAAAAGGGAATAAGG<br>GCGACACGGAAATGTTGAATACTCATACTCTTCCTTTTTCAATATTATTGAAGCA<br>TTTATCAGGGTTATTGTCTCATGAGCGGATACATATTTGAATGTATTTAG |

[a]: The DNA sequences coding for 3’SL (97 nts) and DENV-mini (719 nts) RNAs are colored in red;
[b]: The T7 promoter is colored in green;
[c]: The common upstream sequence targeted by pMVF primer is colored in blue.

**Table S2. The ssDNA primers used for dsDNA template preparations by PCR.**

| Primers | Sequence 5'-3' | Application |
|---|---|---|
| pMVF | GTAACCCACTCGTGCACCCAACTG | Forward primer |
| 689R | AGAACCTGTTGATTCAACAGCACCATTCCA (NaM)TTTCTGGCGTTCTG | Reverse primer for $T_{689}$NaM modification |
| 719R | (NaM)GAACCTGTTGATTCAACAGCACCATTC | Reverse primer for $A_{719}$NaM modification |
| 689/719R | (NaM)GAACCTGTTGATTCAACAGCACCATT CCA(NaM)TTTCTGGCGTTCTGTG | Reverse primer for $T_{689}$NaM /$A_{719}$NaM modification |
| 3'SLR | AGACCCATGGATTTCCCCACACCGG | Reverse primer for native 3'SL and DENV-mini RNAs |

**Table S3. Specific PCR conditions for amplification of the UBP-modified dsDNA templates.**

| Product DNA | 3'SL | | | DENV-mini | | |
|---|---|---|---|---|---|---|
| | $A_{689}$NaM | $T_{719}$NaM | $A_{689}$NaM/ $T_{719}$NaM | $A_{689}$NaM | $T_{719}$NaM | $A_{689}$NaM/ $T_{719}$NaM |
| 2×PCR mix | 25 µL | | | | | |
| Template | 4 ng/µL | 4 ng/µL | 6 ng/µL | 8 ng/µL | 8 ng/µL | 10 ng/µL |
| dTPT3-TP | 1 mM | 1 mM | 1.4 mM | 1 mM | 1 mM | 1.4 mM |
| Forward primer | 0.8 µM | | 1 µM | 0.8 µM | | 1 µM |
| Reverse primer | 0.8 µM | 0.8 µM | 1 µM | 0.8 µM | 0.8 µM | 1 µM |
| dd $H_2O$ | To 50 µL | | | | | |

**Table S4. Typical conditions for *in vitro* transcription of 3'SL and DENV-mini RNAs containing UBP.**

| Product RNA | 3'SL | | | DENV-mini | | |
|---|---|---|---|---|---|---|
| | $A_{689}$TPT3 | $T_{719}$TPT3 | $A_{689}$TPT3/ $T_{719}$TPT3 | $A_{689}$TPT3 | $T_{719}$TPT3 | $A_{689}$TPT3/ $T_{719}$TPT3 |
| 10 ×Transcription buffer | 10 μL | | | | | |
| T7 RNA polymerase | 200 ng/μL | | | | | |
| NaM-modified DNA template | 4 μM | 4 μM | 6 μM | 4 μM | 4 μM | 6 μM |
| rNTP mix | 4 mM each | | | | | |
| rTPT3 or $rTPT3^{A}$ | 0.8 mM | | 1 mM | 0.8 mM | | 1 mM |
| DTT | 10 mM | | | | | |
| DEPC-$H_2O$ | To 100 μL | | | | | |

**Table S5. Overall structural parameters for native, UBP-modified and Nanogold-conjugated 3'SL and DENV-mini RNAs.**

| Sample | $I_0$[a] | $R_g$[a] | $I_0$[b] | $R_g$[b] | $D_{max}$ | MW[c] | MW[d] |
|---|---|---|---|---|---|---|---|
| 3'SL WT | 0.153±0.017 | 35.327±0.582 | 0.158±0.015 | 38.256±0.569 | 129 | 32,100 | 30,229 |
| 3'SL 689-TPT3 | 0.166±0.042 | 35.754±0.233 | 0.179±0.023 | 37.571±0.511 | 127 | 32,300 | 31,898 |
| 3'SL 719-TPT3 | 0.188±0.029 | 35.741±0.900 | 0.193±0.021 | 37.588±0.530 | 127 | 32,300 | 30,910 |
| 3'SL 689/719-TPT3 | 0.165±0.094 | 35.019±0.623 | 0.181±0.097 | 38.580±1.236 | 130 | 32,500 | 30,058 |
| 3'SL 689 Single-Nanogold | 0.189±0.041 | 35.195±1.233 | 0.200±0.055 | 37.280±1.158 | 135 | n/a | n/a |
| 3'SL 719 Single-Nanogold | 0.205±0.019 | 36.304±0.462 | 0.217±0.081 | 40.672±1.210 | 150 | n/a | n/a |
| 3'SL Double-Nanogold | 0.198±0.022 | 39.364±0.518 | 0.209±0.031 | 41.180±1.059 | 156 | n/a | n/a |
| DENV-mini WT | 1.741±0.051 | 84.128±0.187 | 0.339±0.090 | 89.474±1.967 | 309 | 237,270 | 193,173 |
| DENV-mini 689-TPT3 | 1.415±0.065 | 83.257±0.584 | 1.508±0.831 | 90.156±2.504 | 310 | 237,470 | 197,528 |
| DENV-mini 719-TPT3 | 1.633±0.025 | 85.976±0.391 | 1.749±0.018 | 91.656±1.742 | 311 | 237,470 | 210,785 |
| DENV-mini 689/719-TPT3 | 1.589±0.071 | 84.268±0.741 | 1.693±0.065 | 90.527±1.058 | 310 | 237,670 | 205,469 |
| DENV-mini 689 Single-Nanogold | 0.341±0.017 | 88.221±0.546 | 0.361±0.083 | 90.963±1.812 | 313 | n/a | n/a |
| DENV-mini 719 Single-Nanogold | 0.321±0.019 | 87.158±0.618 | 0.338±0.092 | 89.751±1.907 | 310 | n/a | n/a |
| DENV-mini Double- Nanogold | 0.347±0.016 | 89.944±0.852 | 0.360±0.068 | 91.498±1.405 | 316 | n/a | n/a |
| 1.4 nm Nanogold | 0.026±0.018 | 8.095±0.025 | 0.031±0.008 | 8.133±0.058 | 18 | n/a | n/a |

[a] Derived from Guinier fitting; [b] derived from GNOM analysis; [c] MW: molecular weight predicted from sequences; [d] MW: molecular weight calculated based on the power law of volume of correlation; n/a: not applicable.

**Table S6. The coupling efficiencies for Nanogold-conjugated 3'SL and DENV-mini RNAs.**

| Sample | $A_{260}$ | $A_{420}$ | $Con_{Gold}/Con_{RNA}$ | Labeling efficiency |
|---|---|---|---|---|
| 3'SL 689 Single-Nanogold | 2.537 | 0.287 | 0.836 | 0.836 |
| 3'SL 719 Single-Nanogold | 2.602 | 0.305 | 0.864 | 0.864 |
| 3'SL Double- Nanogold | 2.061 | 0.332 | 1.750 | 0.767 |
| DENV-mini 689 Single-Nanogold | 7.536 | 0.119 | 0.738 | 0.738 |
| DENV-mini 719 Single-Nanogold | 6.942 | 0.101 | 0.791 | 0.791 |
| DENV-mini Double- Nanogold | 6.063 | 0.225 | 1.615 | 0.652 |

**Table S7. SAXS data collection parameters and software employed for data analysis.**

| *Data Collection Parameters* | |
|---|---|
| Facilities and parameters | Settings and values |
| Beam line | 12ID-B (APS, ANL) |
| Wavelength (Å) | 0.8857 |
| Detector | Pilatus 1M (SAXS) |
| $q$ range ($Å^{-1}$) | 0.005-0.89 |
| Exposure time (s) | 3 (XSI)-30 (SAXS) |
| Concentration range (mg/ml) | 0.75-3 |
| Temperature (K) | 298 |
| ***Software Employed*** | |
| Primary Data Processing | Matlab/PRIMUS |
| *P*(r) Function | GNOM |

#### Synthetic procedures and characterizations of rTPT3[A]

##### *General*

All solvents and reagents were purchased commercially and used without further purification. For synthetic procedures, all reactions were carried out in oven-dried glassware under an inert atmosphere. Solvents were distilled and/or dried over 4 Å molecular sieves. NMR spectra were recorded on an AVANCE III 1 BAY 400 MHz Bruker NMR spectrometer and the chemical shifts were reported relative to the deuterated NMR solvent used [$^1$H-NMR: CDCl3 (7.26 ppm), DMSO-d6 (2.50 ppm); $^{13}$C-NMR: CDCl3 (77.16 ppm), DMSO-d6 (39.52 ppm)]. Mass spectra were recorded on an Agilent 1200 + G6110A.

##### *Synthetic schemes and procedures*

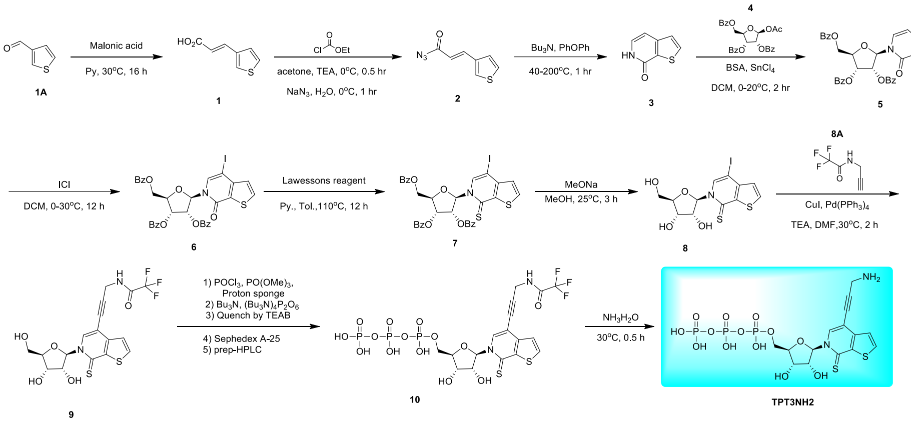


##### *General procedure for preparation of compound 1*

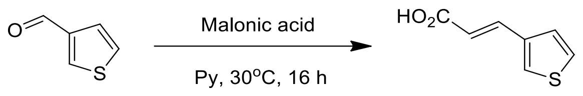


**1A** **1**

To a solution of **1A** (300.0 g, 2.6 mol, 243.9 mL, 1.00 *eq*) in Py (2000 mL) was added Malonic acid (389.7 g, 3.7 mol, 389.7 mL, 1.40 *eq*). The mixture was stirred at 30 °C for 16 hrs. TLC (petroleum ether/ethyl acetate = 1/1, $R_f$ = 0.21) indicated that **1A** was consumed completely and one new spot formed. The reaction was clean according to TLC. Added 12 M HCl adjust pH 2-3, then the solid precipitation, filtered and washed with water, collect filter cake. The filter cake was soluble in EtOAc (5000 mL), added dry $Na_2SO_4$. The organic phase was concentrated under reduced pressure to give a residue. Compound **1** (300 g, crude) was obtained as a light white solid.

**TLC:** petroleum ether/ethyl acetate = 1/1, $R_f$ = 0.21

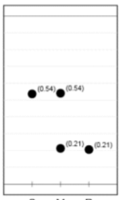


***General procedure for preparation of compound 2***

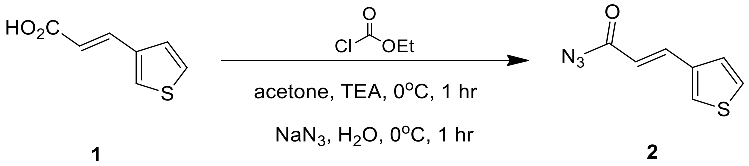


Add acetone (2000 mL) and TEA (236.3 g, 2.3 mol, 324.9 mL, 1.20 *eq*) and compound **1** (300.0 g, 1.9 mol, 1.0 *eq*) to a flask. Add ethyl carbonochloridate (232.3 g, 2.1 mol, 203.7 mL, 1.10 *eq*) dropwise at 0 °C to the mixture. Stir the mixture at 0 °C for 1 hr. TLC (plate 1: petroleum ether/ethyl acetate = 5/1, $R_f$ (material) = 0.05, $R_f$ (product) = 0.40) indicated material was consumed completely. Add $NaN_3$ (151.8 g, 2.3 mol, 1.20 *eq*) in $H_2O$ (750 mL) dropwise at 0 °C to the mixture. Stir the mixture at 0 °C for 1 hr. TLC (plate 2: petroleum ether/ethyl acetate = 5/1, $R_f$ = 0.54) indicated material was consumed completely and one new spot formed. Add $H_2O$ (1.5 L) to the mixture and filtered and check water phase to pH > 9. The filtered cake wash with $H_2O$ (500 mL) and filtered. The filtered cake was dissolved by EtOAc (1.0 L). Wash by brine (1.0 L x 2) and dry with $Na_2SO_4$. Concentrate until remained 1.0 L EtOAc. Add $Ph_2O$ (600 mL) and concentrated to get a solution of product in $Ph_2O$. The crude compound **2** (348.0 g, crude) with white solid in $Ph_2O$ was used into the next step without further purification.

**TLC** (plate 1: petroleum ether/ethyl acetate = 5/1, $R_f$ (material) = 0.05, $R_f$ (product) = 0.40)

**TLC** (plate 2: petroleum ether/ethyl acetate = 5/1, $R_f$ (material) = 0.4, $R_f$ (product) = 0.54)

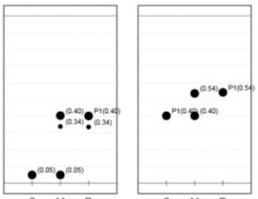


***General procedure for preparation of compound 3***

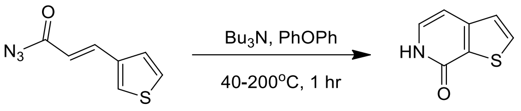


**2** **3**

Add $Ph_2O$ (0.9 L) and $Bu_3N$ (239.8 g, 1.7 mol, 317.2 mL, 3.00 *eq*) to a flask at 150 °C. A heat solution (40-50 °C) of compound **2** (100.0 g, 558.0 mmol, 1.00 *eq*) in $Ph_2O$ (300 mL) was added to the flask at 200 °C. Stir the mixture at 200 °C for 1 hr. TLC (petroleum ether/ethyl acetate = 5/1, $R_f = 0.05$) indicated compound **2** was consumed completely and one new spot formed. The reaction was clean according to TLC. The mixture was cooled to 20 °C and diluted with petroleum ether (10 L). All batches were combined together, the resulting solid was collected by filtration. Combine the two reactions to give a crude product. The filter cake was washed with hexane (5000 mL). The solid obtained was dried under vacuum to give the product. Compound **3** (200.0 g, 1.3 mol, 79.0% yield) was obtained as a yellow solid.

**TLC:** petroleum ether/ethyl acetate = 5/1, $R_f = 0.05$

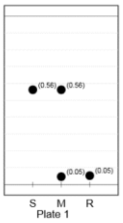


***General procedure for preparation of compound 5***

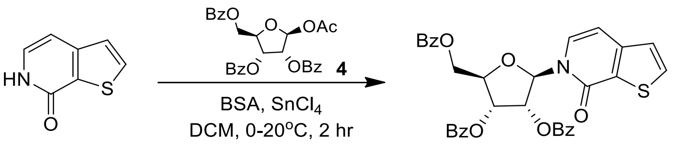


**3** **5**

Added compound **3** (150 g, 992.16 mmol, 1.00 *eq*) and DCM (2500 mL) to a 5 L flask. Then add BSA (242 g, 1.2 mol, 294.3 mL, 1.20 *eq*) to the flask under $N_2$. Stir the mixture for 0.5 hr. Add **4** (550.6 g, 1.10 mol, 1.10 *eq*), then added $SnCl_4$ (310.2 g, 1.2 mol, 139.1 mL, 1.20 *eq*) drop wise to the mixture at 0 °C. Stir the mixture at 20 °C for 1.5 hrs. TLC (petroleum ether/ethyl acetate = 2/1, $R_f = 0.64$) indicated compound **3** was consumed completely and one new spot formed. The reaction was clean according to TLC. Added sat.aq. $NaHCO_3$ (3000 mL) to the mixture and adjust pH 7, then filtered. The filtrate was extracted by DCM (2000 mL x 2). Wash the organic phase with brine (1.5 L) and dry with $Na_2SO_4$, concentrate the organic phase to give crude product. The residue was purified by column chromatography ($SiO_2$, petroleum ether/ethyl acetate = 5/1 to 2/1). Compound **5** (950.0 g, 1.52 mol, 76.4% yield, 95.0% purity) was obtained as a white solid.

**TLC**: petroleum ether/ethyl acetate = 2/1, $R_f = 0.64$

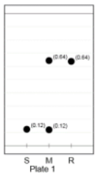


**$^1$H NMR:** 400 MHz, $CDCl_3$. δ ppm 8.17 (d, *J* = 7.2 Hz, 2H), 7.98 (t, *J* = 7.0 Hz, 4H), 7.72 (d, *J* = 5.2 Hz, 1H), 7.45-7.66 (m, 5H), 7.39 (q, *J* = 7.6 Hz, 5H), 7.18 (d, *J* = 5.2 Hz, 1H), 6.79 (d, *J* = 4.8 Hz, 1H), 6.57 (d, *J* = 7.4 Hz, 1H), 6.04 (t, *J* = 5.8 Hz, 1H), 5.88-5.97 (m, 1H), 4.91 (dd, *J*=12.0, 2.8 Hz, 1H), 4.77-4.84 (m, 1H), 4.69-4.76 (m, 1H).

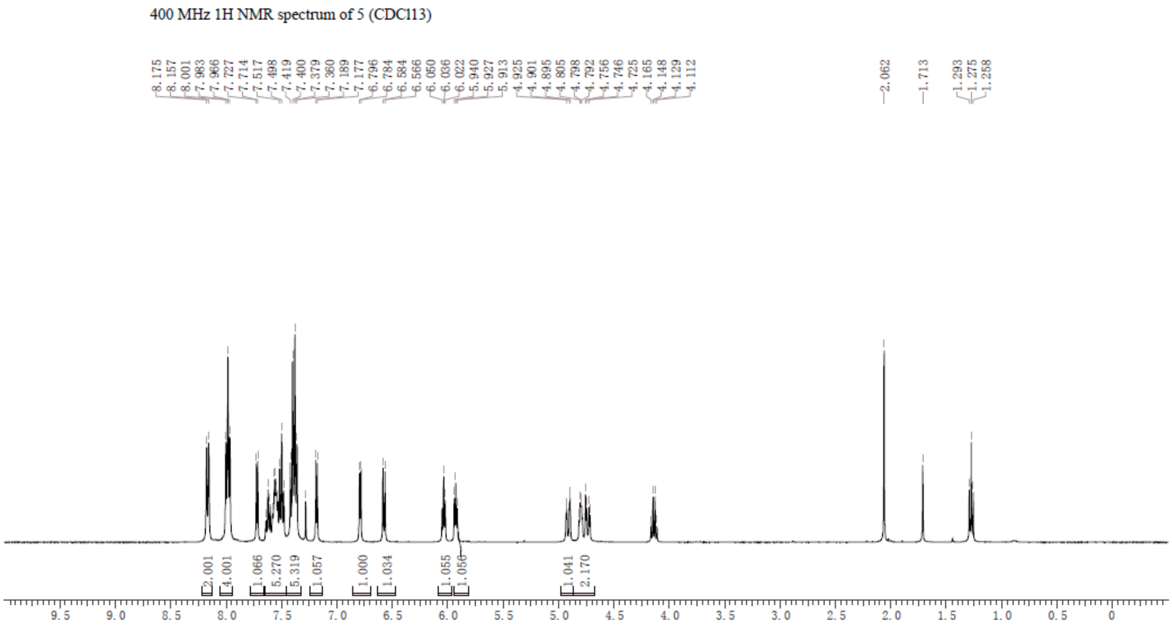


***General procedure for preparation of compound 6***

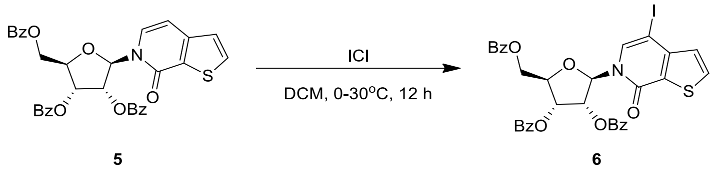


To a solution of compound **5** (340.0 g, 570.8 mmol, 1.00 *eq*)) in DCM (2000 mL) was added ICl (185.4 g, 1.10 mol, 58.3 mL, 2.00 *eq*) at 0 °C. The mixture was stirred at 30 °C for 12 h under exclusion of light. TLC (petroleum ether/ethyl acetate = 3/1, $R_f$ = 0.36) indicated compound **5** was consumed completely and one new spot formed. The reaction was clean according to TLC. The reaction mixture was quenched by addition $Na_2S_2O_3$ (1500 mL) at 20 °C, and then diluted with DCM (2000 mL x 2). The organic layers were concentrated under reduced pressure to give a residue. The residue was purified by column chromatography ($SiO_2$, petroleum ether/ethyl acetate = 5/1 to 3/1). Compound **6** (200 g, 249.5 mmol, 43.7% yield, 90.0% purity) was obtained as a white solid.

**TLC**: petroleum ether/ethyl acetate = 3/1, $R_f = 0.36$

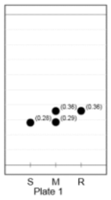

***General procedure for preparation of compound 7***

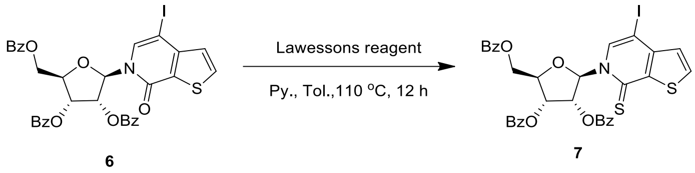

To a solution of compound **6** (140.0 g, 194.04 mmol, 1.00 *eq*) in toluene (1200 mL) was added 2, 4-bis (4-methoxyphenyl)-2, 4-dithioxo-1, 3, 2, 4-dithiadiphosphetane (117.7 g, 291.1 mmol, 1.50 *eq*) and Py (15.4 g, 194.0 mmol, 15.7 mL, 1.00 *eq*). The mixture was stirred at 110 °C for 12 h. TLC (petroleum ether/ethyl acetate = 3/1, $R_f = 0.39$) indicated compound **6** was consumed completely and one new spot formed. The reaction was clean according to TLC. The reaction mixture was concentrated under reduced pressure to give a residue, the residue was diluted with DCM (1500 mL) and extracted with $H_2O$ (1000 mL x 2). The organic layers were concentrated under reduced pressure to give a residue, added 500 mL MeOH stirred for 2 h, filtered and get the cake. The residue was purified by column chromatography ($SiO_2$, petroleum ether/ethyl acetate = 10/1 to 5/1). Compound **7** (100.0 g, crude) was obtained as a yellow solid.

**TLC**: petroleum ether/ethyl acetate = 3/1, $R_f = 0.39$

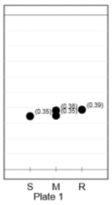

**$^1$H NMR:** 400 MHz, $CDCl_3$, δ ppm 8.07 - 8.13 (m, 3H), 7.97 (dd, *J* = 8.4, 1.2 Hz, 2H), 7.71-7.91 (m, 4H), 7.39-7.59 (m, 5H), 7.23-7.38 (m, 5H), 7.16 (d, *J* = 5.4 Hz, 1H), 5.78-5.87 (m, 2H), 4.88 (dd, *J* = 12.6, 2.4 Hz, 1H), 4.80 (td, *J* = 5.2, 2.6 Hz, 1H), 4.64-4.74 (m, 1H).

400 MHz 1H NMR spectrum of 7 (CDCl13)

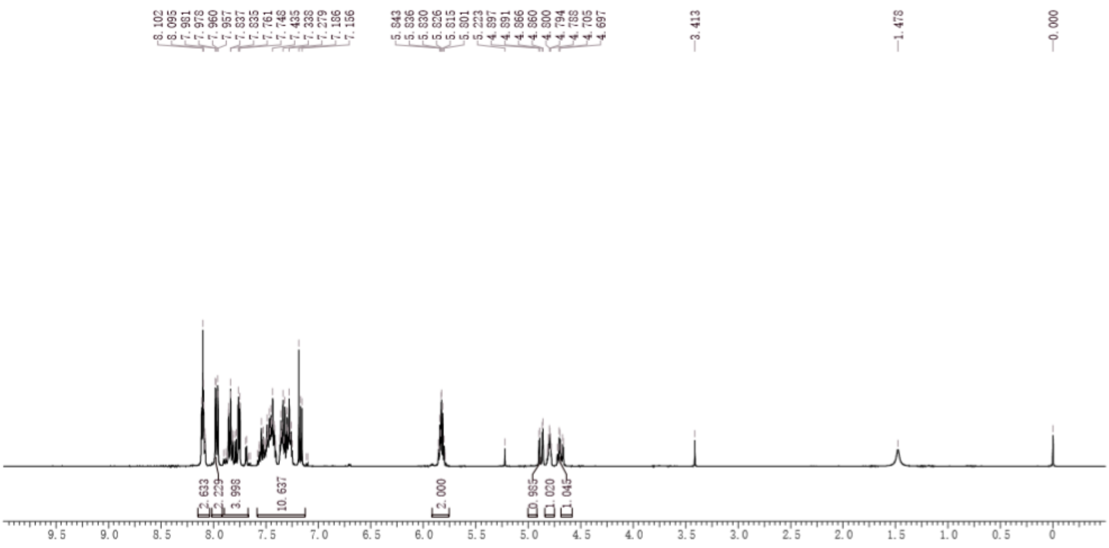

*General procedure for preparation of compound 8*

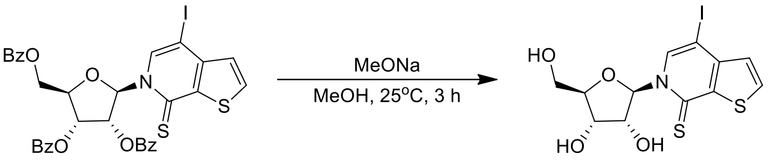


**7** **8**

To a solution of compound **7** (100.0 g, 135.6 mmol, 1.00 *eq*) in MeOH (500 mL) and DCM (500 mL) was added NaOMe (3.66 g, 67.79 mmol, 0.500 *eq*). The mixture was stirred at 25 °C for 3 hrs. TLC indicated compound **7** was consumed completely and one new spot formed. The reaction was clean according to TLC. The reaction mixture was concentrated under reduced pressure to give a residue. The residue was purified by column chromatography ($SiO_2$, DCM/MeOH = 100/1 to 10/1). Compound **8** (40.0 g, 75.3 mmol, 55.5% yield, 80.0% purity) was obtained as a yellow solid.

**TLC**: petroleum ether/ethyl acetate = 3/1, $R_f$ = 0.03

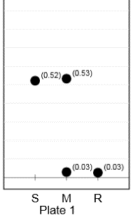


**$^1$H NMR:** ET27024-50-P1a1, 400 MHz, DMSO. δ ppm 9.01 (s, 1H), 8.26 (d, $J$ = 5.4 Hz, 1H), 7.33 (d, $J$ = 5.4 Hz, 1H), 6.81 (d, $J$ = 1.50 Hz, 1H), 5.46-5.66 (m, 2H), 5.12 (d, $J$ = 5.6 Hz, 1 H), 4.00-4.22 (m, 4H), 3.88 (m, 1H), 3.60-3.76 (m, 1H).

400 MHz 1H NMR spectrum of 8 (DMSO)

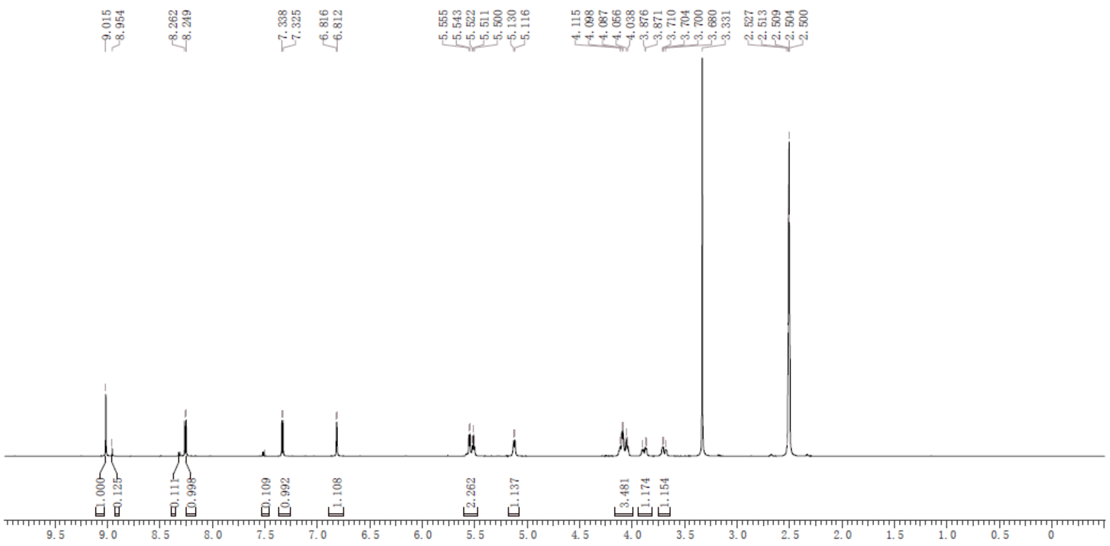

***General procedure for preparation of compound 9***

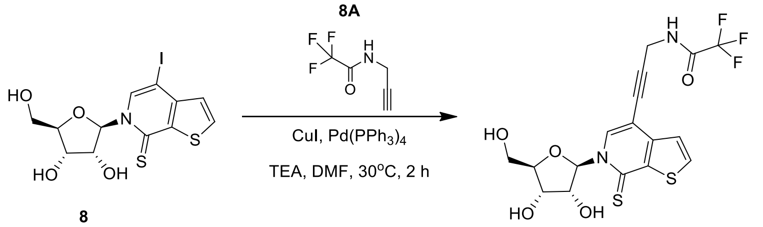


8

9

To a solution of compound **8** (9.0 g, 21.2 mmol, 1.00 *eq*) in DMF (150 mL) was added $Pd(PPh_3)_4$ (2.5 g, 2.10 mmol, 0.100 *eq*) and CuI (806.1 mg, 4.20 mmol, 0.200 *eq*) and TEA (3.20 g, 31.8 mmol, 4.4 mL, 1.50 *eq*), then added **8A** (4.8 g, 31.7 mmol, 1.50 *eq*). The mixture was stirred at 30 °C for 2 h. LC-MS showed compound **8** was consumed completely and one main peak with desired m/z was detected. The reaction mixture was concentrated under reduced pressure to give a residue. The residue was purified by column chromatography ($SiO_2$, DCM/MeOH = 20/1). Compound **9** (9.0 g, 18.1 mmol, 85.4% yield, 90.0% purity) was obtained as a yellow solid.

**LCMS:** (product: RT = 1.054 min, MS cal: 448.4, $[M+1]^+$ = 449.1);

LCMS Chromatogram of 9

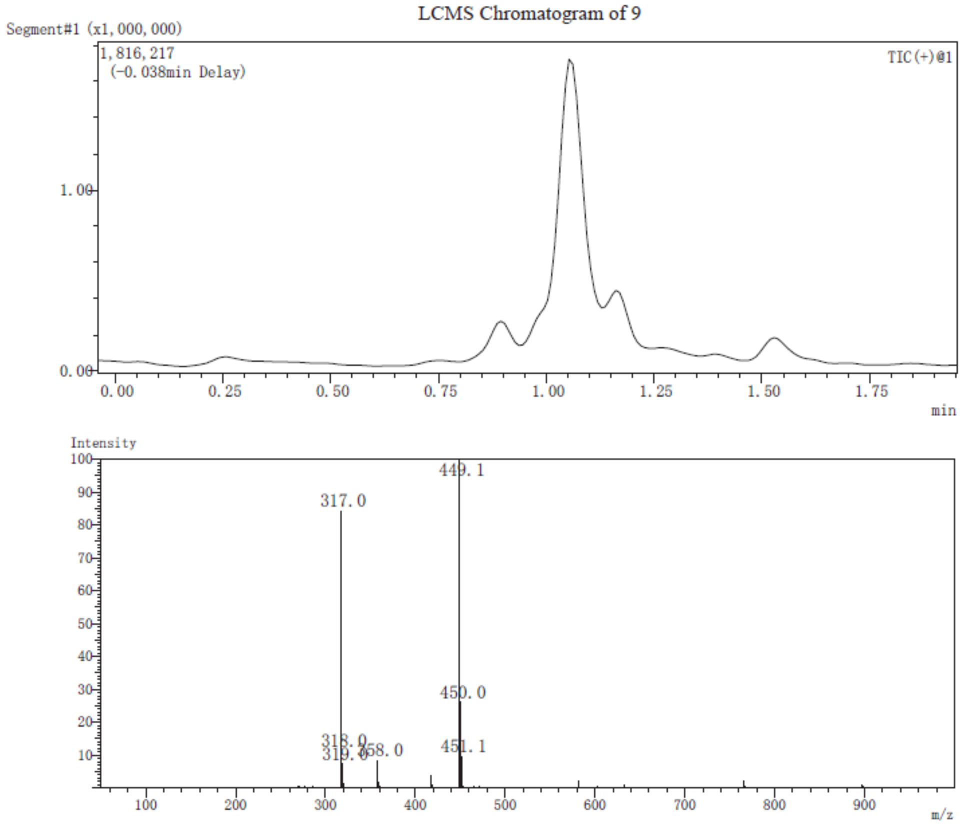


**$^{1}$H NMR:** 400 MHz, MeOD. δ ppm 8.84 (s, 1H), 8.03 (d, $J$ = 5.4 Hz, 1H), 7.46 (d, $J$ = 5.4 Hz, 1H), 6.94 (d, $J$ = 1.4 Hz, 1H), 4.40 (s, 2H), 4.15-4.32 (m, 3H), 4.10 (dd, $J$ = 12.6, 2.0 Hz, 1 H), 3.90 (dd, $J$ = 12.4, 2.2 Hz, 1H).

400 MHz 1H NMR spectrum of 9 (MeOD)

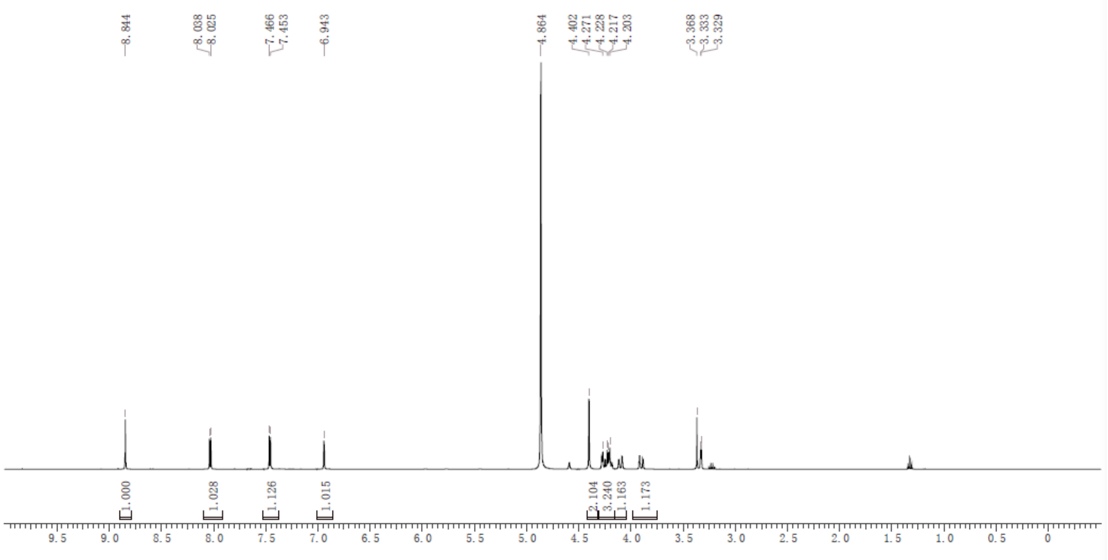


***General procedure for preparation of compound 10***

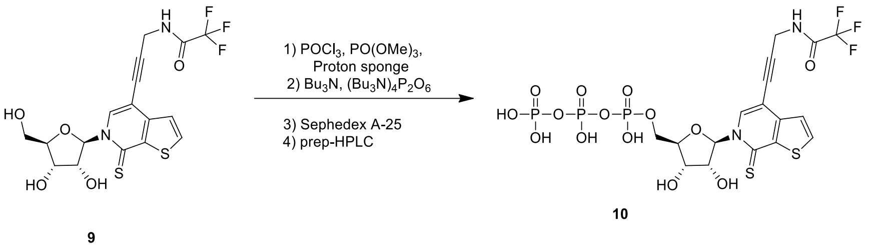


To a solution of compound **9** (2.0 g, 4.5 mmol, 1.0 *eq*) and proton sponge (955.8 mg, 4.46 mmol, 1.00 *eq*) in $PO(OMe)_3$ (20 mL) was added $POCl_3$ (1.00 g, 6.7 mmol, 621.7 uL, 1.50 *eq*). The mixture was stirred at 0 °C for 2 h. LC-MS showed compound **9** was consumed completely and one main peak with desired m/z was detected. The crude product (7.60 g, crude) with yellow oil was used into the next step without further purification.

To a solution of crude product (2.50 g, 4.42 mmol, 1.00 *eq*) in $PO(OMe)_3$ (20 mL) was added N, N - dibutylbutan -1-amine; phosphono dihydrogen phosphate (12.1 g, 22.1 mmol, 5.00 *eq*) and N, N-dibutylbutan-1-amine (4.90 g, 26.5 mmol, 6.3 mL, 6.00 *eq*). The mixture was stirred at 0 °C for 2 h. LC-MS showed material was consumed completely and one peak with desired m/z was detected. Added 1M TEAB to adjust pH7, the reaction mixture was purified by a DEAE Sephadex column (GE Healthcare) with an elution gradient of 0 to 1.0 M TEAB, evaporated to obtain a yellow oil, the crude product **10** (8.0 g, crude) was used into the next step without further purification.

***General procedure for preparation of compound 11(TPT3$^A$)***

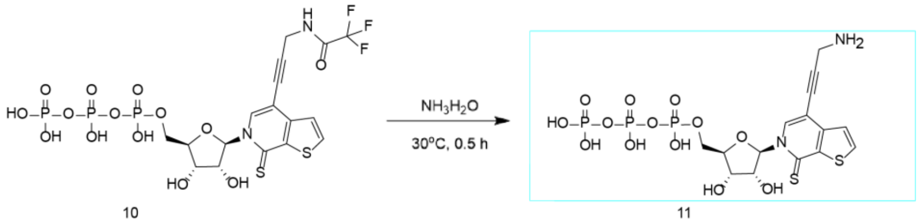


To a solution of compound **10** (5.0 g, 7.3 mmol, 1.00 *eq*) in $NH_3.H_2O$ (5.0 mL) and $H_2O$ (5.0 mL), then stirred at 30 °C for 0.5 h. LC-MS showed compound **10** was consumed completely and one main peak with desired m/z was detected. The mixture of reaction was purified by prep-HPLC (neutral condition, column: Agela Dura Shell C18 250 x 70mm x 10um; mobile phase: [water (10mM $NH_4HCO_3$)-ACN]; B%: 1%-12%, 22 min). **TPT3-NH2 (TPT3$^A$)** (0.85 g, 1.4 mmol, 18.7% yield, 95.0% purity) was obtained as a yellow solid.

**LCMS:** (product: RT = 1.014 min);

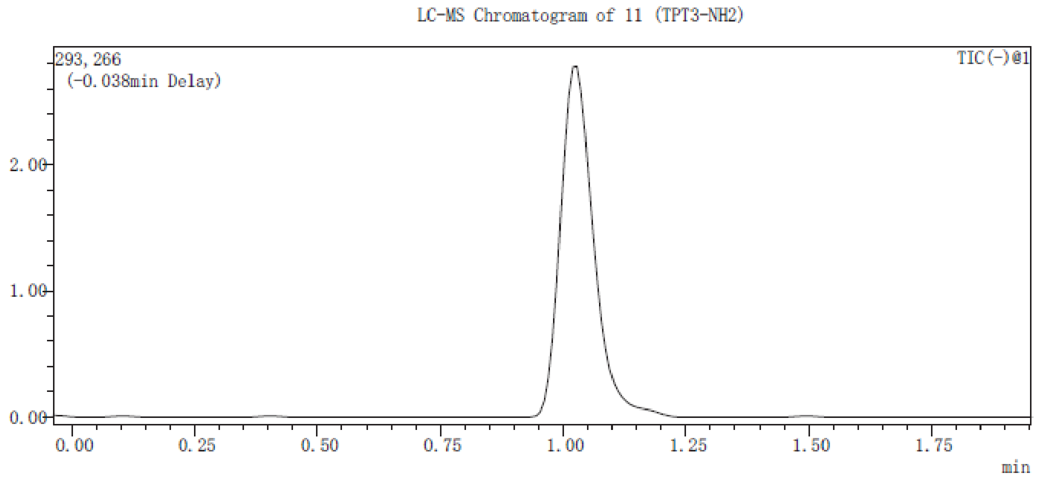


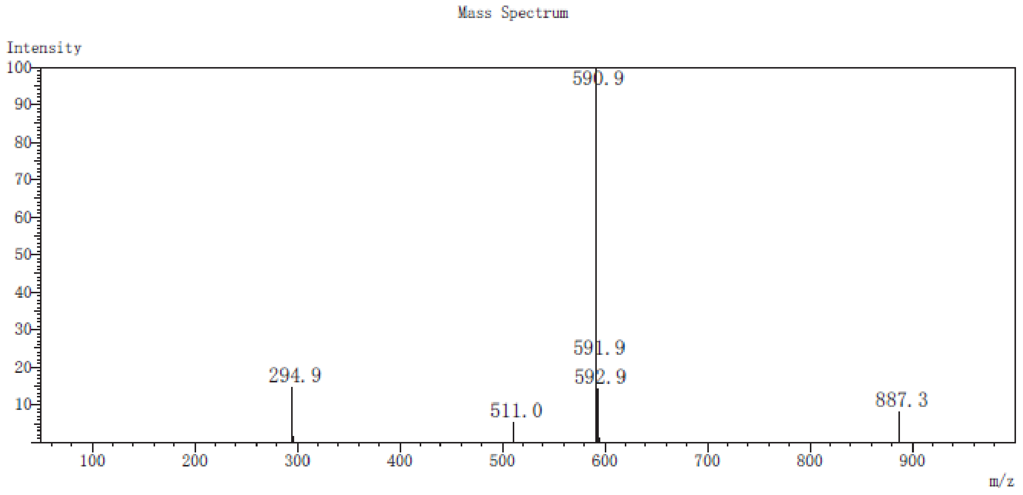


**[1]H NMR:** 400 MHz, $D_2O$. δ ppm 8.49 (s, 1H), 7.90 (m, 1H), 7.24 (m, 1H), 6.68 (s, 1H), 4.47 (m, 1H), 4.20-4.39 (m, 4H), 4.03 (s, 2H).

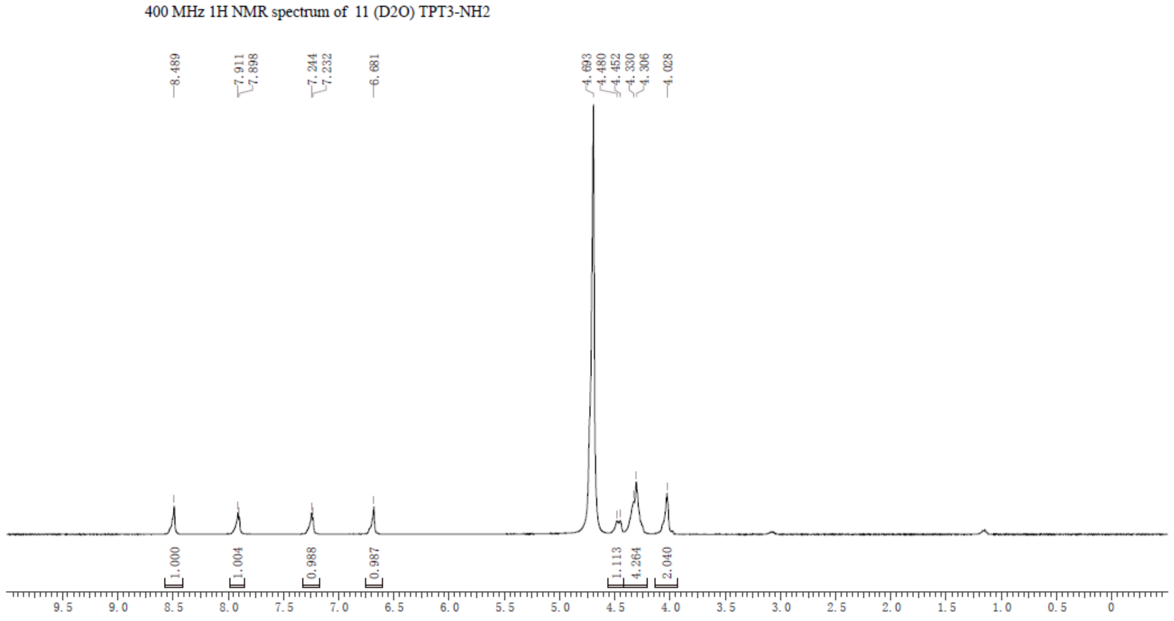


**[31]P NMR:** 400 MHz, $D_2O$. δ ppm -8.27 (d, $J$ = 17.6 Hz, 1P) -11.45 (d, $J$ = 20.6 Hz, 1P) -22.21 (t, $J$ =

19.0 Hz, 1P).

$^{31}P$ NMR spectrum of 11 (TPT3-NH2) (D2O)

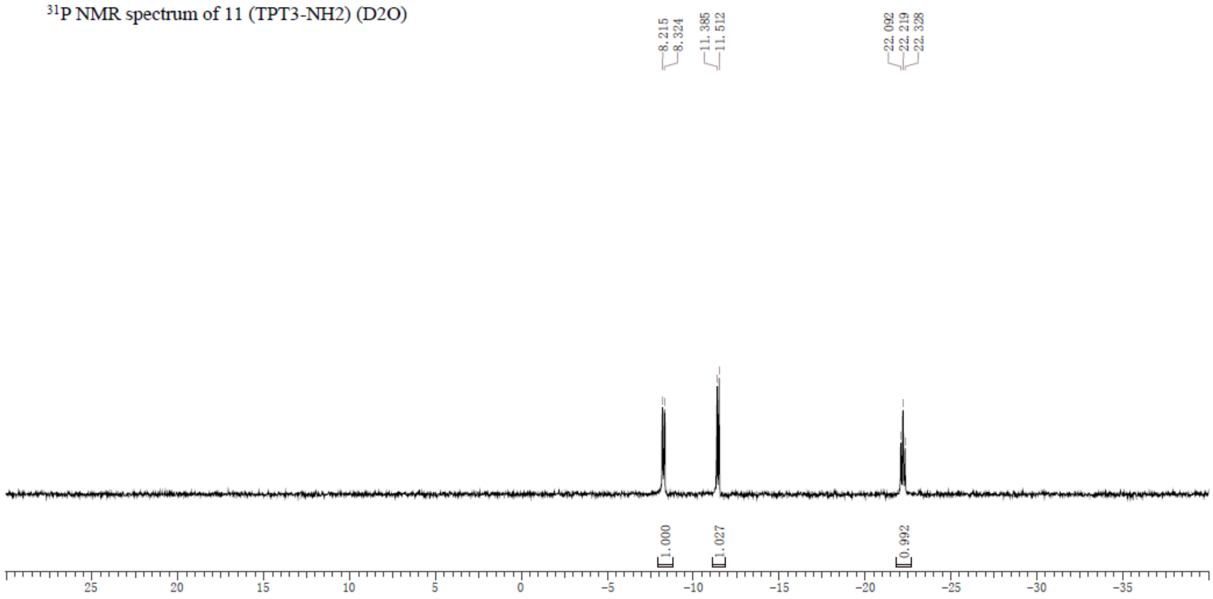

#### References


1. Sztuba-Solinska, J.; Teramoto, T.; Rausch, J. W.; Shapiro, B. A.; Padmanabhan, R.; Le Grice, S. F., Structural complexity of Dengue virus untranslated regions: cis-acting RNA motifs and pseudoknot interactions modulating functionality of the viral genome. *Nucleic Acids Res* **2013,** *41* (9), 5075-89.
